# The tumor suppressor Tip60 inhibits TORC1 signaling in response to microbial acetate to promote autophagy and enterocyte differentiation

**DOI:** 10.64898/2026.07.20.739606

**Authors:** Juliana Batista, Paula I. Watnick

## Abstract

The intestinal microbiota is critical for maintenance of local and systemic immune and metabolic homeostasis in animals, but few molecular mechanisms of action have been delineated. Here, using a *Drosophila* model, we elucidate the role of the microbial fermentation product acetate in maintenance of the intestinal barrier and enterocyte maturation. Tip60 is a lysine acetyl transferase that modifies histone and non-histone targets. We previously showed that Tip60 activates innate immune signaling in enteroendocrine cells in response to microbe-derived acetate. Here we elucidate a distinct mechanism of action in enterocytes. mTOR is a serine-threonine kinase that regulates cell growth and autophagy based on nutrient availability as part of the TORC1 complex.

We report that microbe-derived acetate represses enterocyte TORC1 signaling in a Tip60-dependent manner. This licenses autophagy, which is required to destroy commensal microbes phagocytosed by enterocytes, resulting in bacterial dissemination. Single cell sequencing shows accumulation of poorly differentiated enterocytes in Tip60 knockdown intestines. The microbiota, Tip60, and mTOR have been implicated in the development and progression of colorectal cancer. As accumulation of undifferentiated precursors is a harbinger of malignant transformation and metastasis, we propose our findings provide a mechanistic link between the microbiota, Tip60, and mTOR, epithelial innate immunity and oncogenesis.

## Introduction

The microbes that inhabit the intestines of animals have been linked to obesity, metabolic health, inflammation, and malignancy ^1–10^. Microbial metabolites secreted into the intestine are the principal mediators of these interactions because the intestine has evolved receptors that detect and respond to these metabolites. The common fruit fly, *Drosophila melanogaster*, has been used as an informative, tractable model to understand the genetic basis of these interactions ^11–13^.

The *Drosophila* intestine is comprised of a foregut, midgut, hindgut and rectum (Fig 1A). The foregut includes the crop, a blind pouch that stores ingested contents and the cardia or proventriculus, a bulbus, multilayered structure with a robust immune response ^14,15^. A peritrophic membrane that is synthesized in the proventriculus extends distally to separate the intestinal epithelium from the intestinal lumen with its attendant microbiota ^16^. This membrane is comprised of chitin-binding mucins anchored to an overlying chitin matrix ^14,17–20^. The midgut is separated into three compartments with distinct functions and transcription profiles ^15,21^. The anterior midgut (AMG) expresses genes involved in vitamin, sugar, and protein metabolism. The acidic middle midgut (MMG) likely aids in the digestion of proteins and lipids and blocks passage of commensal and pathogenic microbes into the posterior midgut (PMG) ^22^. The PMG participates in lipid and protein metabolism. As a result of the immune response of the proventriculus and the low pH of the MMG, the PMG is relatively microbe-free ^22^. A pyloric valve separates the PMG from the hindgut and rectum, which are principally involved in water absorption.

**Figure 1:**
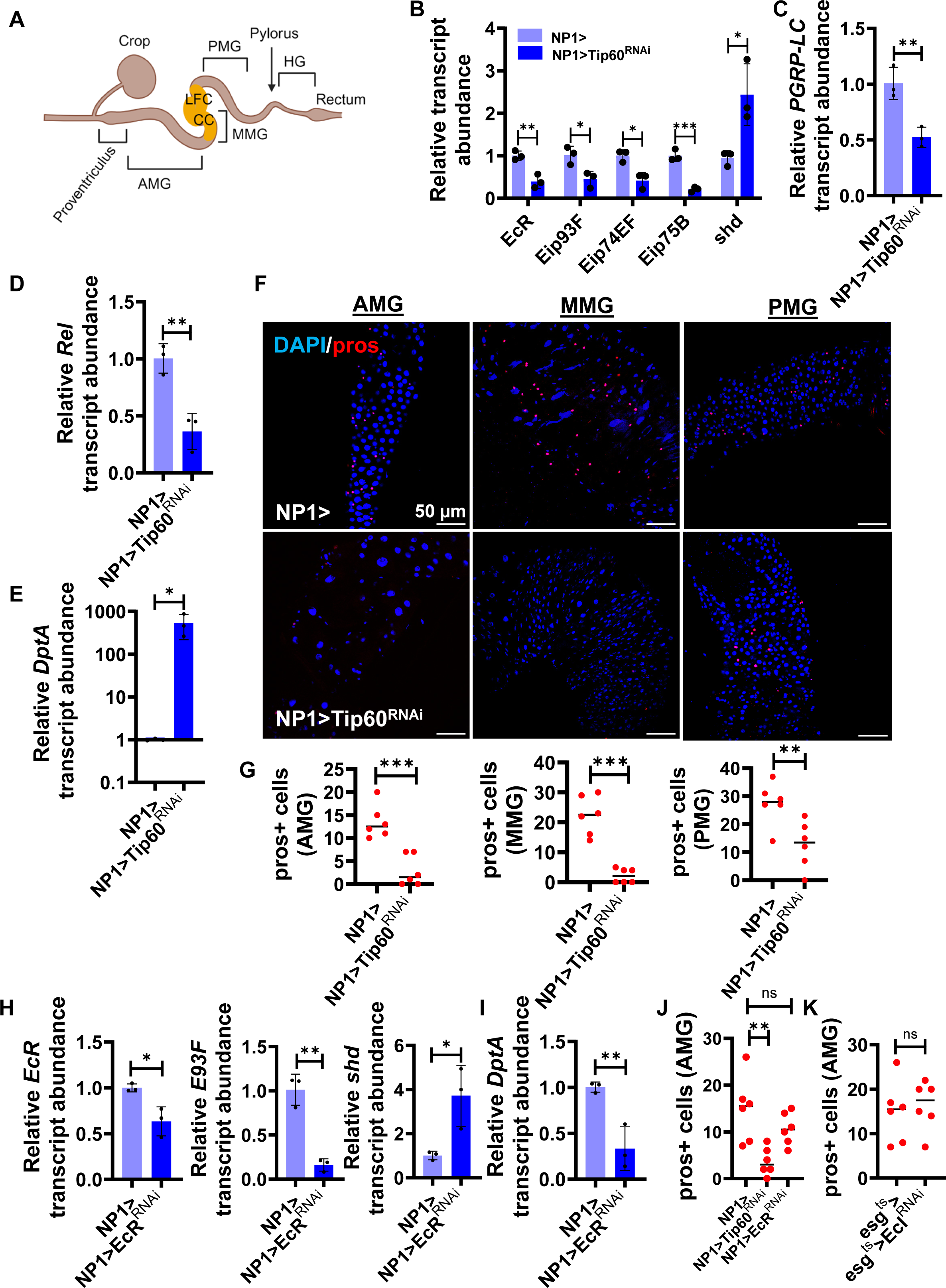
Tip60 regulates 20E signaling in ECs but knockdown of EcR in ECs does not recapitulate the EC Tip60 knockdown phenotype, see also Figs S1 and S2. (A) Compartments of the *Drosophila* intestine. The proventriculus and crop as well as the anterior midgut (AMG), middle midgut (MMG), and posterior midgut (PMG) are shown. Within the MMG, the positions of the copper cells (CC) and the large, flat cells (LF) are shown. qRT-PCR analysis of the indicated (B) ecdysone-regulated and (C-E) IMD pathway-regulated genes in driver-only NP1> and NP1>Tip60^RNAi^ flies. For all qRT-PCR experiments, the mean of biological triplicates is shown. Error bars represent the standard deviation. An unpaired student’s t test was used to assess significance for B-D. A lognormal Welch’s t test was used to assess significance in E. (F) Micrographs and (G) quantification of prospero (pros)+ cells in the AMG, MMG, and PMG of the indicated flies by immunofluorescence. Scale bar 50 µM. The mean of six intestines is shown. A Welch’s t test was used to assess significance. (H and I) qRT PCR of EcR regulated genes in the indicated fly lines. The mean of biological triplicates is shown. Error bars represent the standard deviation. An unpaired t test was used to assess significance. (J) and (K) Enumeration of prospero (pros)+ cells in the intestines of the indicated fly lines. The mean of six intestines is shown. For J, a one-way ordinary ANOVA was used to calculate significance. For K, an unpaired student’s t test was used. **** p<0.0001, *** p<0.001, ** p<0.01, * p<0.05, ns not significant.

The midgut epithelium is comprised of three mature cell types: stem cells, enteroendocrine cells (EEC), and enterocytes (EC) ^13^. Stem cells replenish the epithelium, and complete replacement of the epithelium has been reported to take as little as 4 days or as long as two weeks, depending on the laboratory ^23–25^. This difference may result from the technique used to measure turnover, the growth medium, and/or lab-specific intestinal microbiota.

ECs are the most abundant cell type in the intestine and carry out many essential functions including secretion of proteases and lipases, import of nutrients, peritrophic matrix synthesis, and production of antimicrobial peptides ^21^. The MMG epithelium is comprised of a region of copper cells interspersed with interstitial cells followed by large flat cells, which are specialized ECs ^26,27^. Copper cells are identified by high expression of the developmental transcription factors labial and cut ^27^. Copper cells also express vacuolar ATPases, which are molecular proton pumps that lower the pH of the MMG to less than 3. Large flat cells in the distal MMG are described only by their morphology as their function remains obscure. EEC cells are rare cells that secrete packets of signaling peptides such as tachykinin (Tk) into the hemolymph upon sensing of specific intestinal or systemic signals ^28^. While immature forms of other cell types have not been identified, the unique signature of the enteroblast, an immature cell type that differentiates into ECs, has been defined ^29^.

The *Drosophila* microbiota resides mainly in the AMG ^22^. Although native flies have a more complex intestinal microbiota, that of laboratory flies is simple, comprised mainly of *Lactobacillus* and *Acetobacter* species. Metabolites produced by this community activate the intestinal innate immune response and protect the host during nutritional stress ^30–34^.

The Immunodeficiency or IMD pathway is a TNF-like innate immune signaling pathway that is active both systemically and in all cell types of the *Drosophila* intestinal epithelium ^35–37^. IMD signaling is activated in response to diaminopimelic acid-containing peptidoglycan, which is sensed by the extracellular receptor PGRP-LC and the intracellular receptor PGRP-LE. Through several intermediates, signaling ultimately results in phosphorylation and cleavage of the transcription factor Relish, which translocates to the nucleus to regulate expression of innate immune effectors including antimicrobial peptides (AMPs). Every cell type in the gut expresses components of the IMD pathway, but the spectrum of innate immune effectors produced by each cell type is distinct ^33,38^.

We previously identified microbe-derived acetate as an activator of EEC IMD signaling ^32^. Microbe-derived acetate is imported into Tk+ EEC cells via the EEC cell-specific monocarboxylic acid transporter Tarag and converted into acetyl-CoA. This induces the lysine acetyltransferase Tip60 to activate ecdysone (20E) signaling, which increases expression of PGRP-LC, thereby priming the IMD pathway ^32,39–41^. While Tip60 can acetylate both histone and non-histone targets, in this case, there is evidence that Tip60 acts via acetylation of variant histone 2A ^42–44^.

In these cells, IMD signaling activates expression of the EEC peptides Tachykinin (Tk), Diuretic hormone 31, and neuropeptide F ^33^. Tk, in turn, activates AMP expression and reduces lipid accumulation in ECs through its receptor TkR99D ^32,45,46^.

Because monocarboxylic acid transporters such as Mct1, CG8468, and CG13907 are expressed in all intestinal cell types, we hypothesized that acetate might increase Tip60 activity ECs with a unique outcome ^21,47^. mTOR is a serine/threonine kinase that activates cell growth and represses autophagy in response to nutrient availability as part of the TORC1 complex ^48^. Here we show that, in response to acetate produced by the microbiota, Tip60 in ECs blocks TORC1 signaling. This licenses autophagy, which is required to destroy intestinal microbes that successfully access the EC cytoplasm. Furthermore, single cell sequencing shows that Tip60 knockdown in ECs causes an accumulation of incompletely differentiated ECs with a signature of copper cell precursors. This is accompanied by a loss of mature copper cells and MMG acidification as the adult fly ages. Tip60 controls activation of IMD signaling from EEC cells in response to acetate ^32,33^. Here we identify two additional intestinal innate immune functions regulated by Tip60, namely the destruction of intestinal bacteria that are taken up by ECs or xenophagy and amplification of an acid zone that delays bacterial access to distal regions of the gut. Because all three processes limit the organismal burden and dissemination of commensal and pathogenic bacteria, we propose that intestinal Tip60 plays a central role in intestinal microbial surveillance by limiting bacterial burden in response to the microbial metabolite acetate. The implications of our findings extend beyond innate immunity. Tip60 expression is decreased and mTOR activity is increased in human colorectal cancer, and this correlates with cancer progression and metastasis ^49–51^. Here we find that diminished Tip60 expression results in increased signaling through mTORC1 leading to accumulation of poorly differentiated ECs, which are a prelude to malignancy. Thus, our study provides a mechanistic link between a dysbiotic microbial community, Tip60 and mTOR in the development of colorectal cancers.

## Results

### Knockdown of Tip60 in ECs decreases intestinal 20E signaling but has a phenotype distinct from knockdown of the ecdysone receptor EcR

We previously demonstrated that import of acetate into Tk+ EEC cells enables the lysine acetyltransferase Tip60 to activate 20E signaling ^32^. This function depends on components of the Tip60 complex such as the A isoform of Domino and variant histone 2A. The result is increased expression of PGRP-LC leading to intestinal IMD pathway activation and AMP expression. Because ECs transcribe acetate transporters as well as components of both the 20E and IMD signaling pathways, we questioned whether Tip60 might play a similar role in ECs ^21^. To test this, we knocked down Tip60 in ECs using a Myo1A-Gal4 driver (NP1>). We first measured intestinal expression of genes in the 20E pathway. As shown in Fig 1B, the transcript abundance of the gene encoding *EcR* as well as several genes regulated by 20E signaling were decreased, while that of the 20E synthesis gene *shade* was increased. A similar pattern of expression was detected when Tip60^RNAi^ was expressed in Tk+ EECs ^32^. Expression of the peptidoglycan receptor PGRP-LC, which is indirectly regulated by 20E, was also decreased (Fig 1C) ^41^. IMD signaling controls expression of the NF-KB-like transcription factor Relish and AMPs such as *Diptericin A* (*DptA*). While Tip60^RNAi^ in ECs resulted in decreased Relish expression (Fig 1D), *DptA* expression was increased by several orders of magnitude (Fig 1E).

Tip60 in EECs increases Tk expression in the *Drosophila* AMG. To examine the impact of Tip60^RNAi^ in ECs on Tk expression, we assessed Tk immunofluorescence in the different compartments of the intestine. As shown in Fig S1, Tk+ EECs were decreased in the AMG and MMG but not the PMG of NP1>Tip60^RNAi^ flies. To determine whether this reflected a decrease in total EEC numbers, we assessed staining with the EEC marker prospero (pros) (Fig 1F and G). Tip60^RNAi^ in ECs greatly decreased pros+ cell numbers in the AMG and MMG with a less dramatic effect on the PMG, suggesting that Tip60 in ECs alters differentiation into EECs.

Since Tip60^RNAi^ in ECs decreased the transcript abundance of EcR-regulated genes, we tested whether expression of Tip60^RNAi^ and EcR^RNAi^ in ECs might result in similar phenotypes. EcR^RNAi^ and Tip60^RNAi^ in ECs showed similar patterns of EcR-regulated gene expression, but, in contrast to Tip60^RNAi^, EcR^RNAi^ resulted in decreased expression of the AMP DptA (Fig 1H and I). Furthermore, EC EcR^RNAi^ had no impact on total numbers of EECs (Fig 1J and S2). 20E synthesized in the eclosion muscles of pupa is transported into ISC by the transporter EcI and promotes EEC cell development ^52^. To rule out that, in adult flies, 20E synthesized in response to EC Tip60 might also be transported into ISC by EcI to alter EEC cell development, we expressed EcI^RNAi^ in ISC using the escargot-Gal4 driver (*esg>*) and quantified pros+ cells. As shown in Fig 1K, EcI^RNAi^ in ISCs did not impact EEC numbers. Therefore, while Tip60 regulates 20E signaling and PGRP-LC expression in both Tk+ EEC cells and ECs, this does not explain the phenotype of the NP1>Tip60^RNAi^ intestine.

### EC Tip60^RNAi^ increases intestinal innate immune signaling and expression of cell fate commitment genes

To gain insight into the mechanism of action of Tip60 in ECs, we performed RNA-seq on intestinal samples. Using a threshold of at least a 2-fold change and a p value of 0.05 or less after adjustment for multiple comparisons (padj), 690 genes were significantly differentially regulated with 598 genes increased and 92 decreased (Dataset S1). The Panther overrepresentation test showed that genes in the categories of humoral immune response, mitotic cell cycle process, and cell fate commitment were significantly overrepresented (Dataset S1, sheet 4) ^53^. As shown in Fig 2A, some of the genes most activated by EC Tip60^RNAi^ included AMPs, shown in green, and genes involved in intestinal cell development and differentiation, shown in blue.

**Fig 2:**
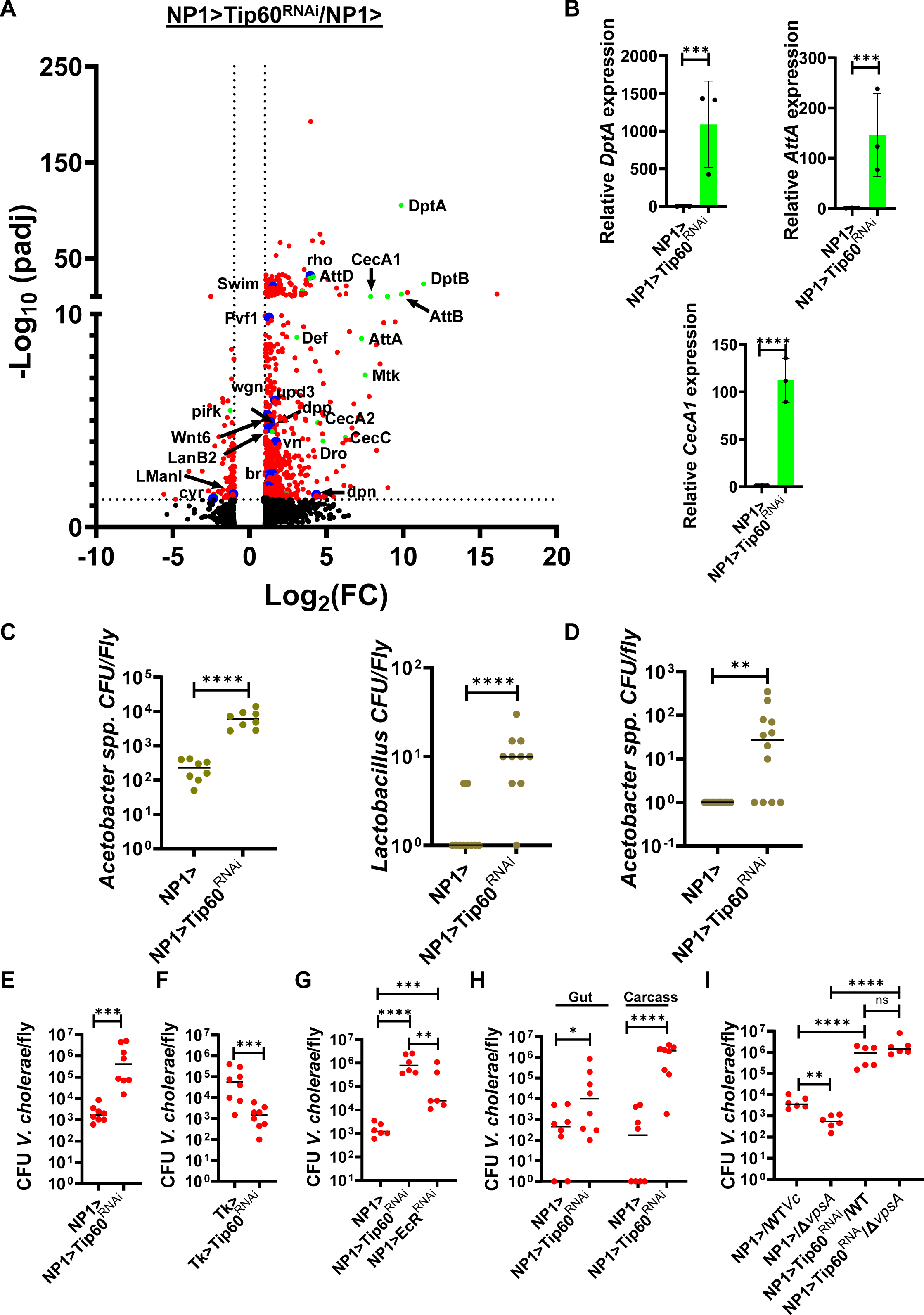
Tip60 knockdown in enterocytes results in activation of the intestinal innate immune response due to microbial penetration of the intestinal epithelium. (A) Volcano plot illustrating the results of an RNA-seq experiment comparing NP1>Tip60^RNAi^ flies with NP1> driver control flies. Points representing innate immune genes are colored green, while points representing cell fate commitment genes are colored blue. Select points are labeled with the gene name. These results represent biological triplicates. Dotted lines indicate a threshold of two-fold change and adjusted p value of 0.05. (B) Transcript abundance of the indicated genes measured by qRT-PCR. The mean of three biological replicates is shown. Error bars indicate the standard deviation. Significance was assessed by applying an unpaired, lognormal t test. (C) Quantification of *Acetobacter* and *Lactobacillus* sp. in flies with the indicated genotypes. The mean of at least 8 whole flies is shown. Statistical significance was assessed using an unpaired, lognormal t test. (D) Quantification of *Acetobacter* sp in the hemolymph of the indicated flies. Each point represents the hemolymph of 10 flies. Twelve groups of flies were analyzed per conditions. A Welch’s lognormal t test was used to calculate significance. (E-G) Quantification of WT *V. cholerae* colonization of the *Drosophila* intestine of flies with the indicated genotypes. The mean of 8 flies is shown. A lognormal Welch’s t test was used to assess significance. (H) Quantification of *V. cholerae* colonization in the gut and carcass of the indicated orally infected flies. The mean of eight flies is shown. An unpaired, lognormal t test data was used to assess significance. (I) Quantification of colonization of the *Drosophila* intestine by WT *V. cholerae* and a Δ*vpsA* mutant that is unable to make a biofilm. The mean burden of eight flies is shown. A lognormal ordinary one-way ANOVA with Tukey’s multiple comparisons test was used to assess significance. **** p<0.0001, *** p<0.001 ** p<0.01, * p<0.05, ns not significant.

### Bacterial dissemination from the intestine is observed with Tip60 knockdown in ECs

We first investigated innate immune signaling. We first measured the intestinal transcript abundance of genes encoding several AMPs and found that these were increased by 2-3 orders of magnitude, supporting our bulk RNA-seq measurements (Fig 2B). We hypothesized that if this were due to direct derepression of the IMD pathway by NP1>Tip60^RNAi^, the intestinal burden of both commensals and pathogens should be decreased. In contrast, if this were a response to inadequate innate immune control of the intestinal microbiota, the microbial burden would be increased. We first measured the burden of the commensal intestinal bacteria *Acetobacter* and *Lactobacillus* in homogenized whole flies (Fig 2C). For both microbes, a several log increase was noted. To assess whether this reflected dissemination of microbes to the hemolymph, we measured the bacterial load of this compartment.

While no bacteria were cultured from the hemolymph of control flies, microbes were detected in the hemolymph of Tip60^RNAi^ flies (Fig 2D). To determine whether the increased bacterial burden was specific to the commensal microbiota, we orally infected flies with the diarrheal pathogen *V. cholerae* for 48 hours, then fed the flies phosphate-buffered saline (PBS) for a 24-hour wash-out period, and finally homogenized flies to measure bacterial load. Again, we observed that the bacterial burden increased by almost 3 logs (Fig 2E). This is the opposite of what was observed for Tip60^RNAi^ in EECs, suggesting a unique role for Tip60 in ECs (Fig 2F) ^32^. In *EcR*^RNAi^ flies, whose intestines express less AMPs rather than more, the burden of *V. cholerae* was increased to a lesser extent (Fig 2G). To test for dissemination of *V. cholerae* beyond the intestine, we separated intestines from the carcass and measured the bacterial load in each. We found that the burden of *V. cholerae* in both the gut and carcass of NP1>Tip60^RNAi^ flies increased (Fig 2H). Techniques that involve dissection or puncture prior to measurement of bacterial burden can be confounded by intestinal laceration. Therefore, we followed up these findings with the following puncture-independent technique. We previously showed that *V. cholerae* forms a biofilm in the posterior region of *Drosophila* intestine ^54^. This biofilm is dependent on the VPS exopolysaccharide synthesized by this bacterium ^55^. When a *vps* synthesis gene such as *vpsA* is mutated, *V. cholerae* is no longer able to form a biofilm, and colonization of the intestine is reduced. We reasoned that, unlike the intestine, *V. cholerae* growth in the hemolymph would not require surface attachment and, therefore, would not be dependent on the *vps* genes. In agreement with our prediction, we found that the Δ*vpsA* mutant had a colonization defect only in control flies but not in Tip60^RNAi^ flies (Fig 2I). Taken together, these data suggest that microbes are able to access the hemolymph from the intestines of NP1>Tip60^RNAi^ but not NP1> flies.

### Knockdown of Tip60 in ECs decreases autophagy

Cells use autophagy to recycle intracellular components such as organelles, proteins, or lipids that are either detrimental to cellular homeostasis, no longer functional, or needed for energy during starvation. This involves fusion of lysosomes with endosomes containing substrates destined for destruction, yielding autophagosomes with low pH and broad degradative capacity ^56^. A specialized form of autophagy termed xenophagy involves the phagocytosis or internalization of extracellular bacteria followed by destruction via the autophagic apparatus ^57,58^. Xenophagy contributes to the barrier function of many types of epithelial cells, which have, therefore, been termed non-professional phagocytes ^59–62^. Our data suggested that AMPs generated by the intestinal epithelium in response to Tip60 knockdown in ECs were not sufficient to prevent dissemination of intestinal bacteria to the hemolymph. We reasoned that xenophagy might be a key innate immune function of the *Drosophila* intestinal epithelium, essential to destroy any microbe reaching the epithelial surface, and hypothesized that Tip60^RNAi^ might block this process. We found support in our RNA-seq dataset. The rare, congenital lysosomal storage disease α-mannosidosis is caused by a deficiency in lysosomal α-mannosidase, which is essential for the lysosomal degradation of N-linked oligosaccharides ^63,64^. In the absence of this enzyme, oligosaccharides accumulate in lysosomes and impede their function. We noted that the transcript abundance of the gene *Lysosomal α-mannosidase* I (LManI) was significantly decreased by Tip60^RNAi^ in our RNA-seq experiment, suggesting an impact on lysosomes (Fig 2A). We confirmed differential regulation of *LManl* in the intestines of Tip60^RNAi^ and Lmanl^RNAi^ flies by qRT-PCR (Fig S3A). We found that Lmanl^RNAi^ in ECs increased *DptA* transcript abundance and increased the burden of commensal bacteria (Fig S3B and C). The similarity of this phenotype to that of NP1>Tip60^RNAi^ flies motivated further exploration of autophagy.

Autophagy is induced by starvation, and defects in autophagy result in increased susceptibility to nutrient deprivation. As a first test, we measured starvation resistance by feeding phosphate-buffered saline (PBS) alone to NP1> and NP1>Tip60^RNAi^ flies as previously described ^46^. As shown in Fig 3A, NP1>Tip60^RNAi^ flies were significantly more susceptible to starvation than NP1> flies. Autophagy-regulated gene 1 (Atg1), a homolog of the mammalian Ulk1, is a kinase that initiates autophagy ^65^. We hypothesized that if the phenotype of NP1>Tip60^RNAi^ flies were the result of a defect in autophagy, then these flies and NP1>Atg1^RNAi^ flies should have similar phenotypes. In fact, the bacterial burden of commensal microbes and *V. cholerae* was similarly increased in NP1>Tip60^RNAi^ and NP1>Atg1^RNAi^ flies, although the NP1>Tip60^RNAi^ phenotype was somewhat more pronounced (Fig 3B and C). To quantify autophagosome formation in the setting of starvation, we starved control NP1>, NP1>Tip60^RNAi^, and NP1>Atg1^RNAi^ 7-day-old flies for 48h and then removed and stained their intestines with lysotracker dye, which fluoresces in response to low pH. Autophagic vesicles were significantly decreased in the intestines of both NP1>Tip60^RNAi^ and NP1>Atg1^RNAi^ flies as compared with NP1> flies (Fig 3D and E). The autophagy protein Atg8a is required for the formation of autophagosomes. To demonstrate that the vesicles observed by lysotracker staining were, in fact, autophagosomes, the intestines of starved flies were stained with an anti-Atg8a antibody. ATG8a immunofluorescence was observed in NP1> flies but not in NP1>Tip60^RNAi^ or NP1>Atg1^RNAi^ flies (Fig S3D and E). Conjugation of Atg8a to the lipid phosphatidylethanolamine (Atg8a-PE) is required for association of Atg8a with the autophagosome ^66^. Atg8a-PE is then degraded with the autophagosome. Conjugated Atg8a-PE is visible as a band that runs faster than Atg8a in a polyacrylamide gel. To determine how Tip60^RNAi^ and Atg1^RNAi^ impacted Atg8a and Atg8a-PE levels, we visualized both forms of Atg8a by Western blot. Under nutrient-replete conditions, Atg8a and Atg8a-PE increased in the intestines of NP1>Tip60^RNAi^ and NP1>Atg1^RNAi^ flies as compared with NP1> controls, consistent with a defect in autophagy (Fig S3F). In starved NP1> flies, levels of Atg8a-PE were greatly decreased, consistent with increased flux through the autophagy pathway. In starved NP1>Atg1^RNAi^ flies, where autophagy initiation is arrested, levels of Atg8a and Atg8a-PE were similar to those in nutrient-replete flies. This shows that starvation has no impact on levels of Atg8a and Atg8a-PE when autophagy is blocked. In nutrient-deprived NP1>Tip60^RNAi^ flies, levels of Atg8a and Atg8a-PE were higher than those in NP1> flies but not as high as those noted in starved NP1>Atg1^RNAi^ flies or fed NP1>Tip60^RNAi^ flies, consistent with a partial response to starvation. Taken together, these results suggest that Tip60 activates autophagy.

**Figure 3:**
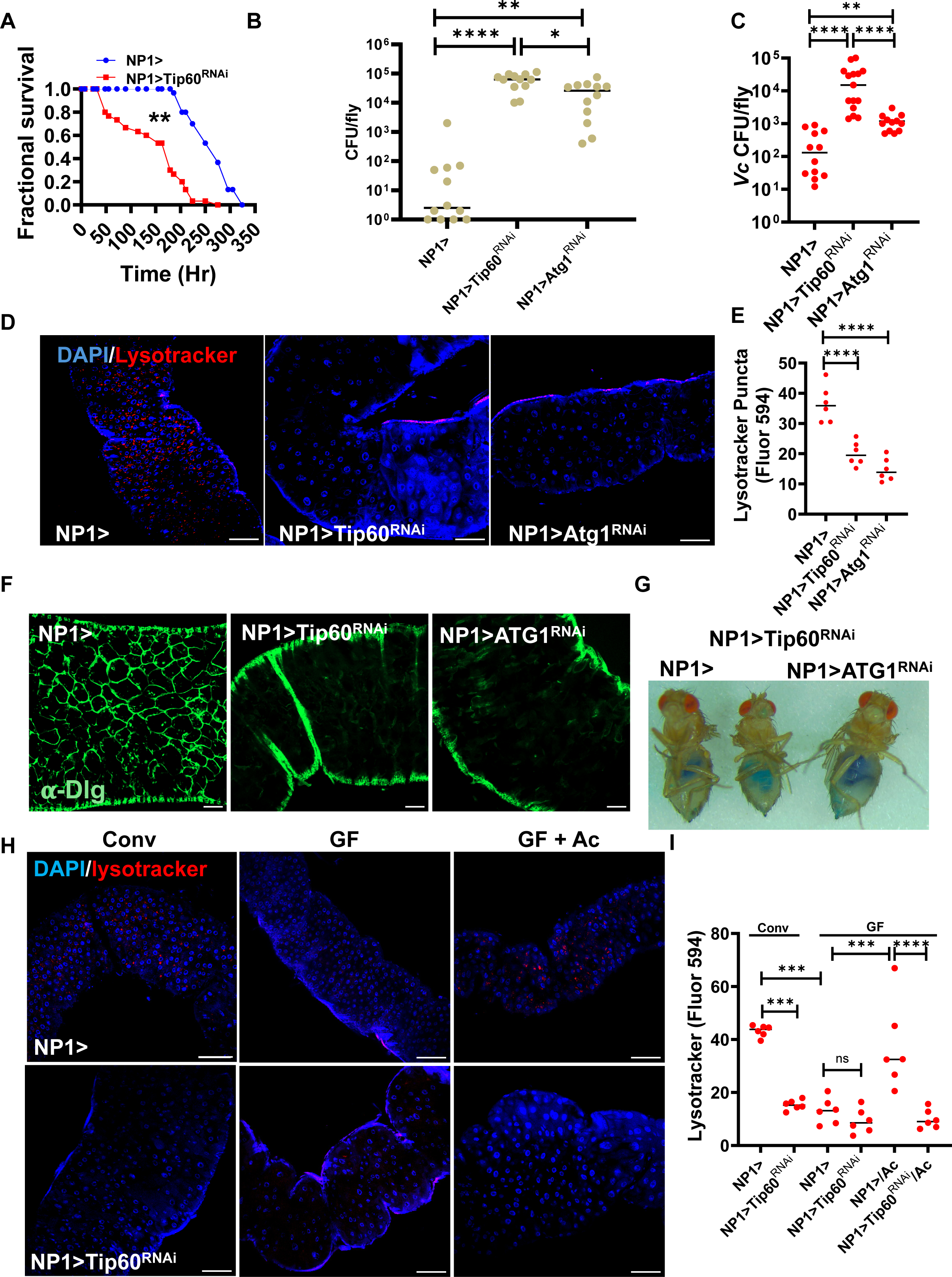
Knockdown of Tip60 in enterocytes inhibits autophagy (see also Fig S3 and S4). (A) Survival of nutrient limitation by the indicated fly lines. Log-rank analysis was used to assess significance. (B) Burden of microbiota in the indicated fly lines. (C) Burden of *V. cholerae* in orally infected NP1> or NP1>Atg1^RNAi^ flies. For B and C, the mean of 12 flies is shown. A lognormal Welch’s one-way ANOVA with Dunnet’s T3 multiple comparisons test was used to assess significance. (D) Imaging and (E) quantification of lysotracker staining in the AMG of the indicated fly lines. The mean of 6 intestines is shown. An ordinary one-way ANOVA was used to assess significance. (F) Dlg1 (α-Dlg) immunofluorescence in the AMG of the indicated fly lines. Scale bar 50 µM. (G) Images of 7 day old flies of the indicated phenotype given access to FD and C blue dye for 48 hours. The dye is contained within the intestine. Lysotracker (H) staining and (I) quantification of the indicated fly lines fed LB alone or treated with antibiotics (GF). Where indicated, LB was also supplemented with 50 mM acetate. Scale bar 50 µm. The mean of at least 6 intestines is shown. An ordinary one-way ANOVA with Tukey’s multiple comparisons test was used to assess significance. **** p<0.0001, *** p<0.001, ** p<0.01, ns not significant.

### Disruption of septate junctions is not the basis of the increased permeability of NP1>Tip60^RNAi^ intestines to microbes

Intestinal septate junctions restrict paracellular access of dye and microbes to the hemolymph ^67–69^. Recently, autophagy in ECs was shown to be essential for maintenance of septate junctions as evidenced by localization of the discs large 1 protein (Dlg) to these junctions. In NP1>Atg1^RNAi^ flies, mislocalization of Dlg was observed as early as 7 days post-eclosion and became more pronounced with age. While paracellular diffusion of an ingested dye was minimal in 10-day old NP1>Atg1^RNAi^ flies, this also increased with age. Because our data suggested that Tip60 in ECs regulates autophagy, we questioned whether permeability of the NP1>Tip60^RNAi^ intestinal epithelial barrier to microbes could be the result of a breakdown in the integrity of septate junctions. Similar to previous reports, we observed mislocalization of Dlg in the intestines of 7-day-old NP1>Atg1^RNAi^ flies (Fig 3F). This was also observed in the intestines of NP1>Tip60^RNAi^ flies but not that of NP1> flies. However, ingested dye was contained within the intestines of control, NP1>Tip60^RNAi^ and NP1>Atg1^RNAi^ flies (Fig 3G and S3G). Because the dye could not cross the intestinal barrier of 7-day-old flies via the paracellular route, it seemed unlikely that bacteria would. To rule out such a scenario, we use a temperature-sensitive driver NP1^ts^> and induced Tip60^RNAi^ expression just four days before imaging. In these flies, a high burden of commensal bacteria was observed despite intact septal junctions visualized by Dlg staining (Fig S4). This supports our contention that bacteria access the hemolymph via the intracellular route in NP1> Tip60^RNAi^ flies. We hypothesize that xenophagy is a component of the intestinal innate immune system that destroys phagocytosed microbes, preventing their access to the hemolymph.

### Microbiota-derived acetate activates autophagy in a Tip60-dependent fashion

In EECs, Tip60 responds to bacterial acetate by activating the innate immune response ^32^. We questioned whether regulation of autophagy by EC Tip60 was also dependent on microbial acetate. To test this, we used an antibiotic cocktail to eliminate the intestinal microbiota of NP1> or NP1>Tip60^RNAi^ flies. We then starved untreated or antibiotic-treated (germ-free) flies for 48 hours. Similar to what was previously observed, lysotracker staining was plentiful in the intestines of starved conventional NP1> flies while very few puncta were seen in the intestines germ-free NP1> flies (Fig 3H and I). Acetate supplementation of germ-free NP1> flies rescued autophagolysosome formation, suggesting that autophagy is increased by this microbial fermentation product. Both conventional and germ-free NP1>Tip60^RNAi^ flies showed no puncta, and acetate treatment of the latter did not increase lysotracker staining. We conclude that activation of autophagy in the setting of starvation is promoted by Tip60 and depends on acetate provided by the microbiota.

### Tip60 activates autophagy by inhibiting signaling through the mTOR pathway

In EEC cells, Tip60 acetylates variant histone 2A to activate 20E signaling ^32^. However, Tip60 also acetylates non-histone targets ^70^. We reasoned that if the phenotype of Tip60^RNAi^ in ECs were the result of a reduction in histone acetylation, the histone deacetylase inhibitor trichostatin should reverse the phenotype. Trichostatin had no effect on *DptA* transcript abundance in NP1>Tip60^RNAi^ intestines (Fig 4A). We then considered that Tip60 might regulate autophagy via a non-histone target. The *Drosophila* serine/threonine kinase mTOR participates in the multi-protein complex, mTORC1, which activates cell growth and survival in response to nutritional signals by repressing autophagy and increasing protein translation via phosphorylation of the ribosomal protein kinase S6K and the inhibitor of protein translation initiation 4EBP (Fig 4B) ^71,72^. Interestingly, mTOR and Atg8a have recently been identified as members of the *Drosophila* Tip60 acetylome, suggesting that Tip60 could either regulate autophagy directly or via mTOR ^73^. Phosphorylation of the ribosomal protein S6 kinase (S6K) is commonly used to detect signaling through the mTORC1 complex. We, therefore, used antibodies against pS6K in a Western blot of the intestines of starved *Drosophila* to determine whether the TORC1 complex was activated by NP1>Tip60^RNAi^ (Fig 4B). In fact, pS6K was increased in NP1>Tip60^RNAi^ intestines. As expected, inhibition of autophagy by NP1>Atg1^RNAi^ had no impact on pS6K levels. The small molecule rapamycin, in complex with FKBP12, binds directly to mTOR to inhibit mTORC1 function ^74^. Levels of pS6K decreased in the intestines of all genetic backgrounds when flies were fed rapamycin. Lipidated Atg8a and Ref(2)p, a homolog of mammalian p62 that recruits ubiquitinated proteins to the autophagosome for digestion, are markers of autophagy.

**Figure 4.**
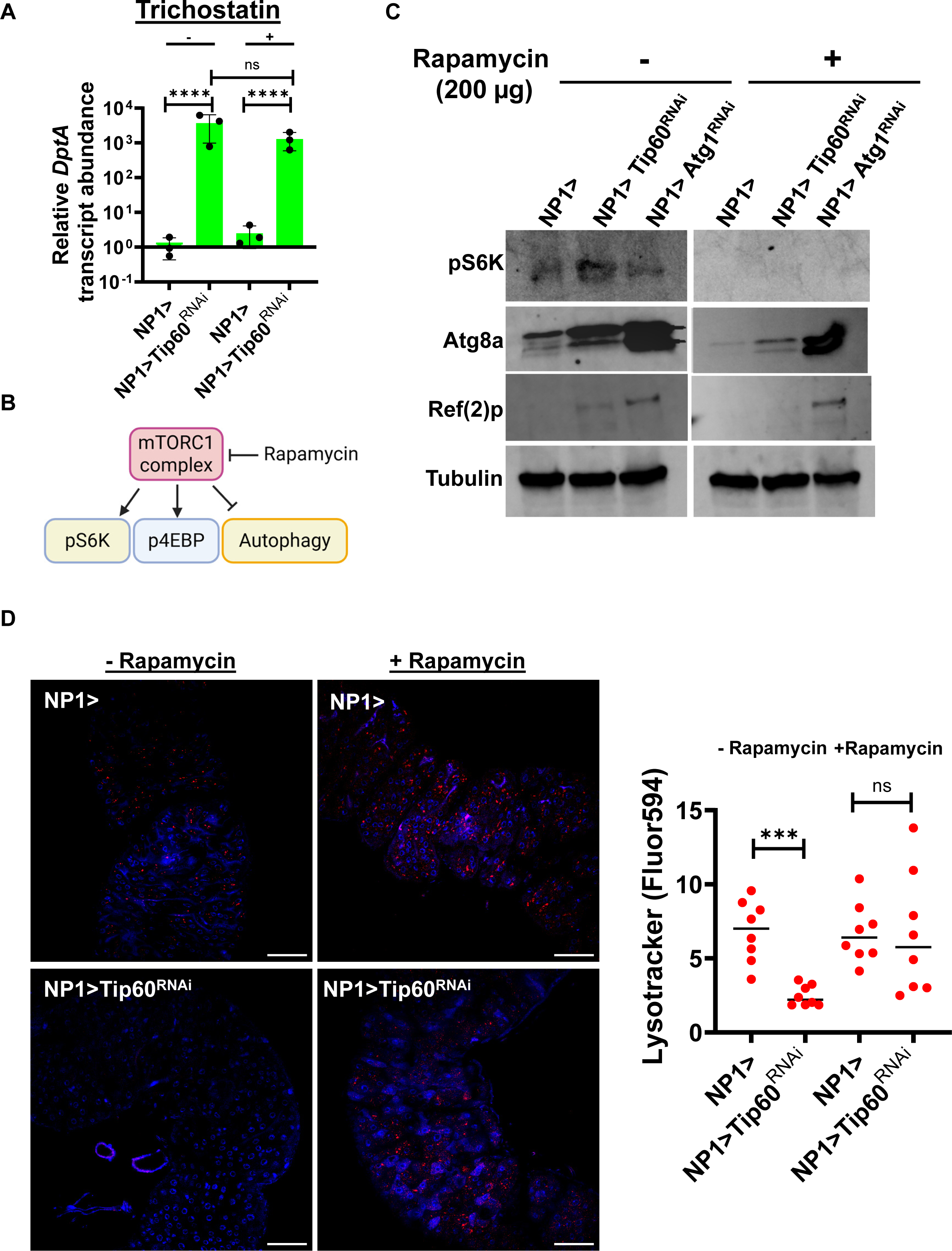
Tip60 acts upstream of Atg1 and autophagy initiation. (A) qRT-PCR analysis of DptA in the intestines of NP1> and NP1>*Tip60*^RNAi^ flies fed the the histone deacetylase inhibitor trichostatin. The mean of three biological replicates is shown. A lognormal one-way ordinary ANOVA with Tukey’s multiple comparisons test was used to assess significance. (B) Diagram showing the components of the pathway connecting mTorc1 to autophagy. Created in BioRender. Watnick, P. (2026) https://BioRender.com/3lv1bq9. (C) Western analysis of the indicated proteins in the indicated fly lines without (-) or with (+) oral administration of 200 µg/ml rapamycin to flies. (D) Lysotracker staining and quantification in the AMG of NP1> and NP1>Tip60^RNAi^ flies with or without rapamycin treatment. The mean of six intestines is shown. An unpaired t test was used to assess significance. **** p<0.0001, *** p<0.001, ** p<0.01.

Therefore, we assessed levels of these species in the intestines of nutrient-deprived NP1>, NP1>Tip60^RNAi^ and NP1>Atg1^RNAi^ flies. Ref(2)p and unlipidated Atg8a were increased in the intestines of starved NP1>Atg1^RNAi^ and NP1>Tip60^RNAi^ flies, suggesting inhibition of autophagy initiation. We reasoned that if Tip60 activated autophagy by directly acetylating a component of the autophagy machinery, oral administration of rapamycin would have no effect on autophagy in the NP1>Tip60^RNAi^ intestine. In contrast, if Tip60 inhibited autophagy by acetylating and inhibiting a component of the mTOR signaling pathway, rapamycin should rescue the NP1> Tip60^RNAi^ phenotype. In fact, rapamycin rescued autophagy in NP1>Tip60^RNAi^ flies as evidenced by decreased levels of Atg8a and Ref(2)p and increased lysotracker staining (Fig 4B and C). In contrast, rapamycin failed to reverse the increased levels of Atg8a and Ref(2)p in the intestines of NP1>Atg1^RNAi^ flies. These data support our hypothesis that Tip60 acts upstream of Atg1, and at or above the level of the TORC1 complex.

### Tip60 regulates systemic and intestinal development

Both mTor signaling and autophagy regulate development ^69,75,76^. We first explored the effect of Tip60 knockdown in ECs on systemic development. NP1>Tip60^RNAi^ flies were approximately 16% shorter and emerged more slowly than NP1> pupae (Fig 5A-C). In addition, many of the pupae either died before or became stuck during emergence (Fig 5D). Adult NP1>Tip60^RNAi^ flies were smaller and weighed less than NP1> flies (Fig 5E and F). The intestines of NP1>Tip60^RNAi^ flies were also noted to be small and dysmorphic (Fig 5G and H). Previous work suggests that Tip60 regulates 20E signaling cell autonomously ^32^. To rule out an effect of 20E signaling in ECs on gut size, we compared the intestines of NP1>Tip60^RNAi^ and NP1>EcR^RNAi^ flies. As shown in Fig 5G and H, the intestines of NP1>EcR^RNAi^ flies were similar to those of controls. This indicates that intestinal Tip60 plays an important role in development that is independent of 20E signaling.

**Fig 5:**
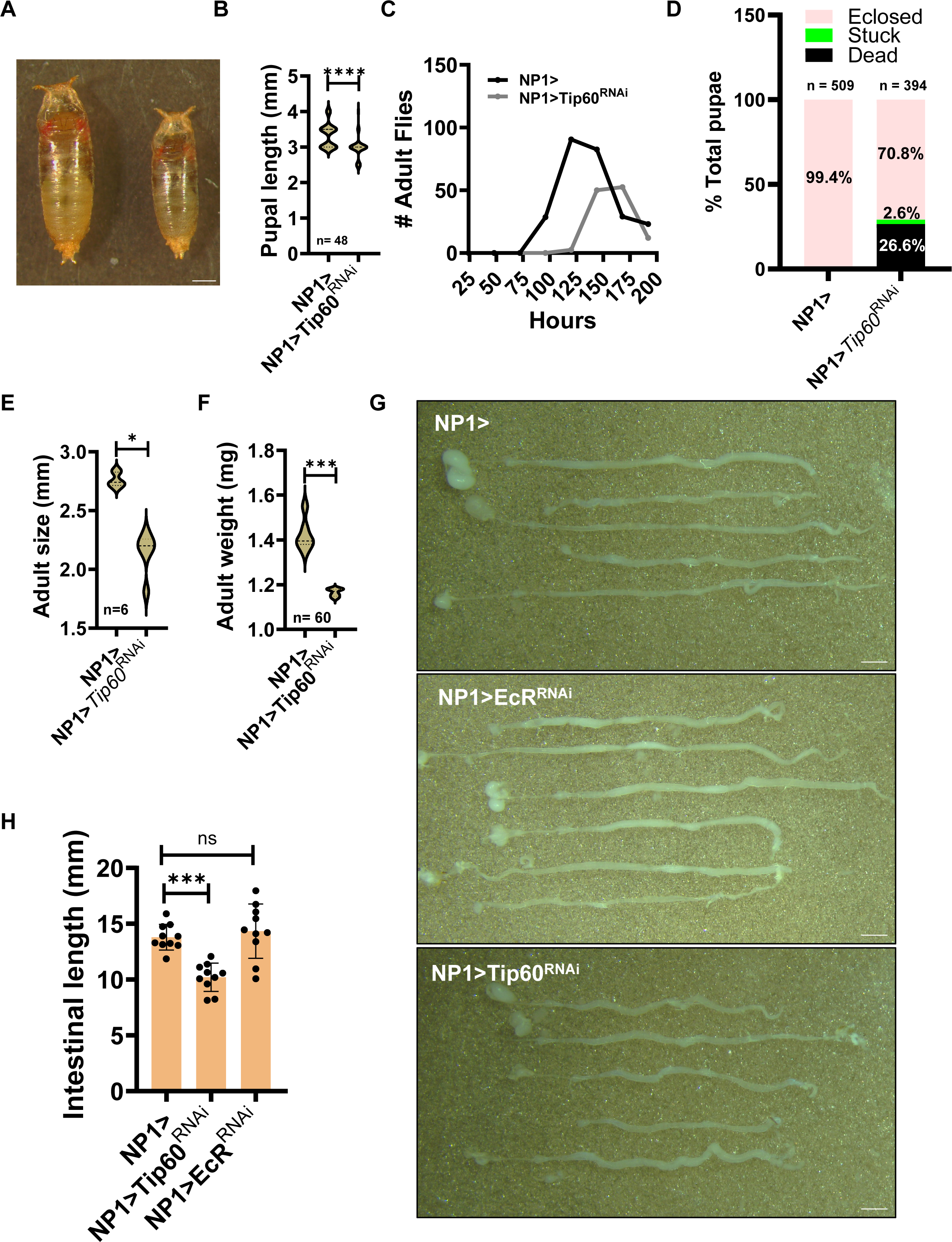
Enterocyte Tip60 is essential for normal *Drosophila* development. (A) Image and (B) quantification of length of pupae formed by NP1> driver control and NP1>*Tip60*^RNAi^ flies. 48 pupae were evaluated. A student’s t test was used to assess significance. Measure bar 1 mm. (C) Numbers of eclosed flies over time for the indicated fly lines. (D) Percent of eclosed adults, adults unable to emerge from pupal sacs (stuck), and dead pupae. The number of flies evaluated is indicated as n. (C) Quantification of adult fly size and weight. A Welch’s t test was used to assess significance. (E and F) Size and weight of adult flies of the indicated genotype. For (E), 6 flies were evaluated. For (F), six groups of 10 flies were evaluated. A Welch’s t test was used to assess significance. (G) Images of the intestines of the indicated fly lines. Measure bar 1 mm. (H) Length of the intestines of the indicated fly lines. The mean of ten intestines is shown. An ordinary one-way ANOVA was used to assess significance. **** p<0.0001, *** p<0.001, * p<0.05, ns not significant.

### Autophagy activated by Tip60 is essential for copper cell replenishment in the adult *Drosophila* intestine

The dysmorphic intestines of NP1>Tip60^RNAi^ flies and decreased EECs suggested aberrant intestinal development. To better characterize this, we carried out single cell sequencing (sc-seq) on the intestines of 7-10 day old NP1> and NP1>Tip60^RNAi^ flies. Cell assignments were based on a previous SC-seq study of the *Drosophila* midgut ^77^. Different numbers of immature and mature cell types to be present in the intestines of control and knockdown flies (Fig 6A and B, Dataset S2). In agreement with our previous immunofluorescence data, NP1>Tip60^RNAi^ intestines had fewer EECs. Equal numbers of mature AMG and PMG ECs were observed in NP1> and NP1> Tip60^RNAi^ flies. In contrast, in NP1>Tip60^RNAi^ intestines, cells in the copper cell cluster were increased 3-fold, while large flat cells were decreased approximately 2-fold, suggesting an impact of Tip60^RNAi^ on MMG development.

**Fig 6:**
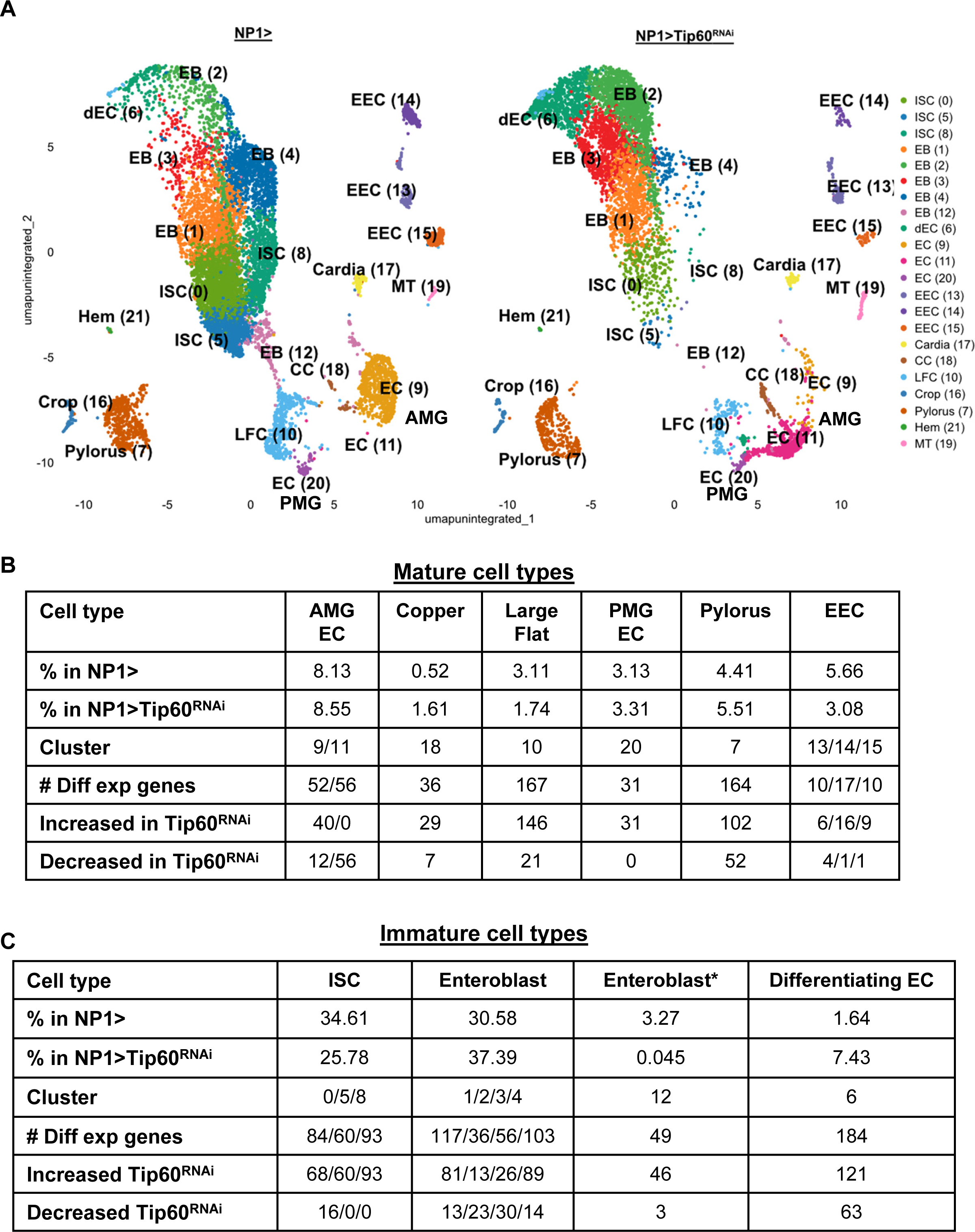
Single cell sequencing suggests that Tip60^RNAi^ in enterocytes retards enterocyte maturation, see also Fig S5. (A) Umap representing sc-seq-based clustering of different cell types in the intestines of the indicated fly lines. ISC intestinal stem cells, EB enteroblasts, dEC differentiating enterocytes, EC enterocytes, EEC enteroendocrine cells, CC copper cells, LFC large flat cells, Hem hemocytes, MT microtubules. Numbers in parentheses indicate the cluster. Table of statistics for (B) the indicated mature cell types and (C) the indicated immature cell types. Cluster indicates the clusters that comprise the cell type, % indicates the percent of the total comprised by the indicated cell type. # Diff exp genes is the number of differentially expressed genes for each cluster in each cell type. The number of genes whose transcription is increased or decreased for Tip60^RNAi^ is given below.

Development and regeneration of the copper cell region is controlled by decapentaplegic (dpp), an ortholog of vertebrate bone morphogenetic proteins 2 and 4 ^78,79^ (Fig S5A). Dpp binds to the thickveins (tkv) receptor to activate transcription of the homeodomain transcription factor, Defective proventriculus (dve). Dve activates expression of the transcription regulator labial (lab), marking these dve+lab+ cells as copper cell precursors. Lab repression of dve results in the differentiation of these precursors into mature copper cells, which are dve-lab+, while cells that remain dve+ and do not express lab are termed interstitial cells ^27,80^. To determine whether copper cell differentiation was impaired by Tip60^RNAi^, we used our sc-seq data set to assess expression of *lab* and *dve* in copper cells. As shown in Fig S5B, lab expression was present in both NP1> and NP1>Tip60^RNAi^ copper cells. While very little dve expression was present in the copper cell cluster of NP1> intestines, significant expression remained in that of NP1>Tip60^RNAi^ intestines (Fig S5C). To further examine this, we enumerated copper cells by immunofluorescence using an anti-lab antibody. As shown in Fig 7A, at 3 days post-eclosion, there were similar numbers of copper cells in the intestines of NP1>, NP1>Tip60^RNAi^, and NP1>Atg1^RNAi^ flies. However, at 10 days post-eclosion, very few copper cells remained in the intestines of NP1>Tip60^RNAi^, and NP1>Atg1^RNAi^ flies. When copper cells are functional, the MMG is characterized by the ability to generate a luminal pH of 2-3. To test the functionality of copper cells, we fed 7-10 day old NP1> and NP1>Tip60^RNAi^ flies the pH-sensitive dye bromophenol blue, which is yellow at pH<3 and blue at pH 7, dissected and imaged the intestines, and quantified the area of the acidic region. As shown in Figs 7B and C and Fig S6, the alkalinity of the NP1>Atg1^RNAi^ and NP1>Tip60^RNAi^ middle midguts were much less than that of NP1> flies. We conclude that autophagy, which we have shown is activated by Tip60 and microbe-derived acetate, is essential for copper cell differentiation in the adult intestine.

**Fig 7:**
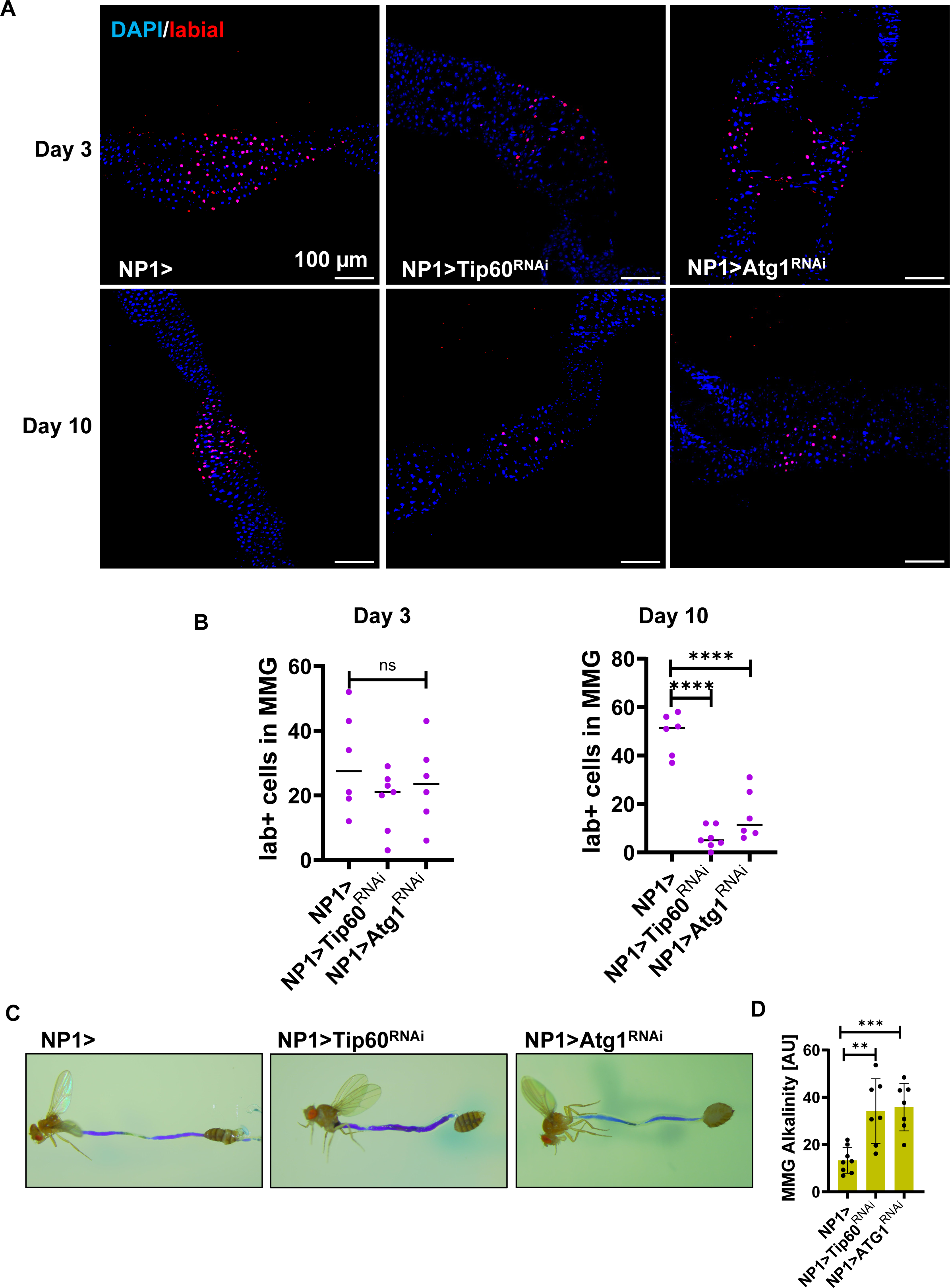
Tip60 is essential for differentiation of copper cells in the adult intestine, see also Fig S6. (A) lab staining and (B) quantification of lab+ cells in the MMG of NP1>, NP1>Tip60, NP1>Atg1 flies. The mean of six intestines is shown. A one-way ordinary ANOVA with Dunnett’s multiple comparisons test was used to assess significance. (C) Images of intestines of the indicated fly lines fed 0.1% bromophenol blue for 24 hours. (D) Relative alkalinity of the intestines of the indicated flies. The mean of at least 7 intestines is shown. A one-way ordinary ANOVA with Dunnett’s multiple comparisons test was used to assess significance. **** p<0.0001, *** p<0.001, ** p<0.01, ns not significant.

### Evidence that differentiating enterocytes in the intestines of NP1>Tip60^RNAi^ flies are copper cell precursors

The intestines of NP1>Tip60^RNAi^ flies had approximately 4.5-fold more differentiating ECs, suggesting a block in differentiation. Because mature copper cells were decreased in the MMG of 10-day old NP1>Tip60^RNAi^ flies, we hypothesized that these differentiating ECs might be MMG precursors. This cell cluster had 184 significantly differentially regulated genes, more than any other cluster. To garner support for our hypothesis, we examined the differentially regulated genes in this cluster. This cell cluster had 184 significantly differentially regulated genes, more than any other cluster. Of these,121 were increased, while 63 were decreased. Of the 100 genes whose transcript abundance was most increased in NP1>Tip60^RNAi^ intestines, 15 were exclusively highly expressed in the MMG (Table S1) ^21^. Of these 15, 7 were uniquely upregulated in the differentiating EC cluster, seven were also differentially expressed in a subset of ISC and EB clusters, and two were additionally differentially expressed in the copper cell cluster (cluster 18). Differential regulation of the copper transporter 1B (Ctr1B), which is essential for copper uptake into copper cells, was also unique to this cell cluster and upregulated approximately 10-fold. This supports our hypothesis that the accumulation of differentiating enterocytes in the MMG of adult NP1>Tip60^RNAi^ flies is indicative of a block in copper cell differentiation that leads to MMG dysfunction.

## Discussion

The widely conserved Tip60 lysine acetyltransferase participates in DNA damage repair, chromatin remodeling, and transcription regulation by acetylating both histones and non-histone proteins such as the ataxia-telangectasia protein kinase, the serine/threonine kinase Bub1, and aurora B kinase ^81–85^. In *Drosophila melanogaster*, Tip60 plays critical roles in oogenesis and activation of the EEC cell innate immune response in the intestine ^32,43^. Here we show that Tip60 in ECs of the *Drosophila* intestine activates autophagy in response to microbial acetate by inhibiting TORC1 signaling. This both prevents intestinal microbes from accessing the hemolymph and maintains the microbicidal acidity of the MMG by promoting differentiation of copper cells. Based on these findings and our previous report that Tip60 activates IMD signaling through EECs, we propose that Tip60 is a master, multi-cell type regulator of the intestinal innate immune response to the microbial product of fermentation acetate ^32^.

Autophagy is known to play a role in cellular differentiation in *Drosophila* through the degradation of transcription factors ^75,86,87^. Here we show that production of acetate by intestinal microbes contributes to normal copper cell differentiation by activating autophagy. Copper cell differentiation requires a decrease in dve expression, which is activated by dpp. Our single cell sequencing data suggest that dve expression is not appropriately extinguished in NP1>Tip60^RNAi^ copper cells, which suggests continued dpp activity. In *Drosophila*, endosomes act as a sink for extracellular dpp by internalizing this ligand bound to its receptor Tkv ^88,89^. While some proportion of this reservoir can be re-externalized when needed, the endosomal contents may also be degraded by autophagy ^90–92^. We hypothesize that when autophagy is diminished by Tip60 blockade, dpp signaling cannot be extinguished, dve expression persists and undifferentiated copper cell precursors accumulate.

We present evidence that Tip60 disrupts mTORC1 signaling to activate autophagy in response to the microbial metabolite acetate. In mammals, modulation of the mTOR pathway and autophagy by lysine acetylation has been shown, but no lysine acetylase or protein target has been identified. For instance, one report showed that HDAC inhibitors block mTOR signaling and increase autophagy in cancer cells ^93^. Another showed that the HDAC Sirtuin 1 increases autophagy by inactivating signaling through the mTOR pathway ^94^. Last, a study showed that microbiota-derived butyrate, a known HDAC inhibitor, induced apoptosis in cancer cells by decreasing mTOR signaling ^49^. Many of these studies aim to treat cancer, obesity, and aging through targeting of lysine acetylation. Therefore, a more complete understanding of the role of protein acetylation in manipulation of the mTOR pathway including the identity and selectivity of the lysine acetylase is essential. The identification of Tip60 as a lysine acetylase that impedes signaling through mTOR and increases autophagy is critical step in a lysine acetylase-targeted therapy for cancer.

Poorly differentiated cells are a harbinger of malignant transformation in human colorectal cancer. Furthermore, sepsis caused by a single intestinal bacterium, most often *Clostridium septicum* or *Streptococcus bovis*, can be a presenting feature of disease ^95^. Here we find that decreased expression of Tip60 in enterocytes results in a similar constellation of symptoms, namely, poorly differentiated intestinal cells and dissemination of intestinal bacteria to the hemolymph. This suggests that a defect in autophagy may be responsible for the connection between monobacterial sepsis and asymptomatic intestinal malignancy.

Tip60 expression is downregulated in tissues obtained from many types of cancers including lung, breast, gastric, and colorectal malignancies and expression levels are correlated with tumor progression ^50,96–100^. The mechanisms by which Tip60 has been shown to influence malignant transformation and metastasis in experimental systems are as diverse as its cellular targets. These include DNA its role in DNA repair, cell cycle regulation, and cellular differentiation^101–103^. Deregulation of the mTOR pathway has similarly been implicated in oncogenesis. The mTOR signaling pathway activates cell growth and inhibits autophagy to promote tumorigenesis, and inhibitors of mTOR are currently used in chemotherapeutic regimens ^48,74,104^. While reports of an interaction between the mTOR pathway and Tip60 exist, they are conflicting and have not elucidated an underlying mechanism ^105–107^. The significance of our findings lies in forging a link between the intestinal microbiota, Tip60, the mTOR pathway, and autophagy, all of which have all been conclusively linked to oncogenesis. We propose that we have uncovered an elusive molecular mechanism that connects intestinal dysbiosis to oncogenesis and provide a simple genetic model to explore the pathophysiology of this complicated process.

## Acknowledgements

This work was supported by NIH R01AI158247 to P.I.W. Anti-TK antibodies were generously provided by Jan Veenstra. The NP1-Gal4 (Myo1A-Gal4) driver flies were kind gifts from Norbert Perrimon. Stocks obtained from the Bloomington Drosophila Stock Center (NIH P40OD018537) were used in this study. Microscopy images were acquired at the Microscopy Resources on the North Quad (MicRoN) core at Harvard Medical School. We thank Paola Montero Lopis at the MicRoN core for providing expertise with image acquisition and quantification.

## Author contributions

J.B. and P. I. W. designed the experiments. J. B. performed the experiments. J. B. and P. I. W. analyzed the data. J.B. and P.I.W. wrote the manuscript. Both authors reviewed, edited, and approved the manuscript.

## Star methods

### Experimental Model

#### Bacterial strains and growth conditions

For colonization experiments, the wild-type *Vibrio cholerae* strain 01 El Tor C6706, strain 2, and the corresponding biofilm deficient Δ*vpsA* strain were cultivated in LB broth (Miller formulation, Difco) overnight at 27°C and then diluted into fresh LB broth as described below ^54,108,109^. To assess intestinal colonization with *V. cholerae*, homogenate dilutions were spotted onto LB agar (Difco) supplemented with streptomycin (100 µg/mL, Difco) and incubated at 27°C for 24 hours. For quantification of the commensal microbiota, homogenate dilutions were spotted onto deMan, Rogosa and Sharpe (MRS) agar (Millipore) and incubated at 27°C for 36 hours.

#### Drosophila strains and husbandry

Fly stocks were cultured on standard fly food containing 16.5 g/L yeast, 9.5 g/L soy flour, 71 g/L cornmeal, 5.5 g/L agar, 5.5 g/L malt, 7.5% corn syrup, and 0.4% propionic acid in a humidified incubator on a 12-hour day-night cycle at 25°C. 3-10 day old female flies were used in all experiments. For experiments with the temperature-sensitive driver NP1^ts^>, flies were raised at 18 degrees and moved to 29 degrees for 3 days prior to dissection and imaging. For generation of microbe-depleted animals, 3-10 days old female flies were placed in vials containing fly food with an antibiotic cocktail consisting of ampicillin (500 μg/mL), tetracycline (50μg/mL), rifampicin (400 μg/mL), and Streptomycin (100 μg/mL) for 4 days. For oral trichostatin treatment, at least 10 female flies per condition were placed in vials containing cellulose acetate plugs infused with LB alone or supplemented with 10μM trichostatin (TSA; Cayman, 58880-19-6).

Treatment was continued for 3 days prior to harvesting of the intestine for qRT-PCR. For experiments with rapamycin, flies were starved for 48 h and then placed in vials with PBS containing 1.5%agar and rapamycin 200 μg/ml (Thermo Fisher) for 3 days prior to removal of the intestine. The following fly stocks were supplied by the Bloomington Drosophila Stock Center: TRiP (BL 36303), Tip60^RNAi^ (BL 28563), Ecr^RNAi^ (BL 58286), ATG1^RNAi^ (BL 26731), LmanI^RNAi^(BL 44473) and DomA^RNAi^ (BL 65873). The Tk-Gal4, NP1-Gal4, NP1-Gal80 and esg-Gal4 lines were kindly provided by the Perrimon lab.

### Method Details

#### RT-qPCR and RNA-seq

Total RNA was extracted from at least 10 fly intestines using TRIzol reagent (Thermo Fisher Scientific 15596026), isolated with the Direct-zol RNA MiniPrep Plus kit (Zymo Research R2070), following the manufacturer’s instructions, and quantified using a NanoDrop 1000 spectrophotometer. For RT-qPCR, 500ng of total RNA was used to synthesize cDNA using the iScript cDNA Synthesis Kit (Bio-Rad 1708891). iTaq^TM^ Universal SYBR® Green supermix (Bio-rad 1725121) and a QuantStudio ^TM^ Real-time PCR system 3 (Thermo Fisher Scientific) were used to quantify mRNA targets. Relative expression was calculated using the comparative C_T_ method normalized to rp49. The primers used for the RT-qPCR reactions are listed in Data set S4. For the RNA-seq, library preparation and sequencing was performed by the Molecular Biology Core Facilities (MBCF) at the Dana Farber Cancer Institute (DFCI).

#### Quantification of intestinal commensal and *V. cholerae* numbers

For assessment of commensal burden in whole flies, groups of 6-12 3-10 day-old flies per genotype raised on standard fly food were collected and place in a 96 deep well plate prepared with a 4.5 mm glass bead and 100 μl of phosphate buffered saline (PBS). Flies were then homogenized for two minutes in a TissueLyzer III system (Qiagen). For assessment of *V. cholerae* burden, overnight *V. cholerae* culture were diluted 10-fold in fresh LB broth, and 3mL of the bacterial suspension were infused into cellulose plugs placed at the bottom of a standard fly vial. Twenty flies were placed in the vial for 48 hours. At this point, 12 randomly chosen flies per fly line were placed in individual compartments of a 96 deep well plate and homogenized as described above. Dilutions of homogenates were prepared, and commensal or *V. cholerae* burden was assessed by plating. For quantification of bacterial burden in the hemolymph, the abdomens of 10 flies per group were pierced with a fine needle. Flies were then placed in a 600 μL microcentrifuge tube with the bottom tip removed. This tube was inserted into a 1.5 mL microcentrifuge tube and centrifuged for 5 minutes at 5,000 rpm. The hemolymph that collected at the bottom of the 1.5 mL tube after centrifugation was suspended with 100 μL pf PBS, serially diluted, and plated onto MRS agar plates.

#### Immunofluorescence

The intestines of 10 female flies (3-10 days old) per genotype were dissected and fixed in 4% formaldehyde (check lot)/PBS solution for at least 20 minutes at room temperature. After fixation, the midguts were washed three times with 0.1% PBT (PBS supplemented with Tween 20) and then incubated in blocking buffer (PBT + 0.1% Triton X-100 + 2% BSA) at room temperature for 1 hour after which primary antibodies were added and the incubation was continued overnight at 4 °C. The dissected midguts were incubated overnight with mouse anti-prospero (1:500, Abcam Ab196361), rabbit anti-GABARAP (anti-Atg8a) (1:500, Cell Signaling 13733), mouse anti-dlg (DSHB AB_528203), guinea pig anti-labial (1:500, Gift from Dr. Benjamin Ohlstein), rabbit anti-Tk (1:500, Gift from Dr. Jan Veentra). After washing three times with PBT, the samples were incubated with fluorophore-conjugated secondary antibodies anti-rabbit Alexa Flour 594 conjugate (1:1000, Invitrogen A11012) anti-guinea pig Alexa Flour 594 conjugate (1:1000, Invitrogen A11076), anti-mouse Alexa Flour 488 conjugate (1:1000, Invitrogen A11001), and 4′,6-diamidino-2-phenylindole (1 μg/ml, DAPI) for one hour at room temperature. All incubations were performed in blocking buffer. After washing with 0.1 Triton X-100/PBS, the intestines were mounted in Vectashield mounting medium. Images were collected using a Zeiss LSM 980 confocal microscope using a 20X or 40X objective.

#### Smurf assays

Seven day old flies were placed in a vial containing standard fly food with 2.5% FD and C blue dye for 48 hours. Flies were imaged using a SMZ800N stereo microscope and a Digital Sight 1000 microscope camera.

#### Starvation survival assays

Three groups of ten flies each per genotype were placed and maintained in vials containing cellulose acetate plug infused with 1x PBS. The number of viable flies was recorded twice a day until all flies were dead.

#### Western Blot Analysis

Twenty 3–10-day-old female flies were homogenized as described for bacterial counts. Fifty μl of 6x Laemmli buffer was added, and samples were heated at 95°C for 10 minutes. After centrifugation to pellet debris, the supernatant was removed and 15 μg of total protein was used to load the 4-20% polyacrylamide gel (Bio-rad) and transferred to a PVDF membrane over 7 minutes using a Trans-blot Tubo (Bio-rad) with the settings 1,3A, 25V. Membranes were incubated overnight with rabbit anti-pS6K (1:500, R&D systems), rabbit anti-GABARAP (anti-Atg8a) (1:1000, Cell Signaling), anti-rabbit anti-Ref2P (1:1000, Abcam), and rabbit anti-tubulin (1:1000, Cell signaling 2144). The secondary antibodies used were anti-rabbit IRDYE 680RD (LicorBio 926-68071) and anti-mouse IRDYE 800CW (LicorBio 926-32210).

#### Developmental assays

To evaluate the number of eclosed adult flies, all crosses were started on the same day, and the number of newly eclosed adult flies was quantified twice daily. To measure length, 8-day-old pupae were imaged with a metric ruler using a SMZ800N stereo microscope and a Digital Sight 1000 microscope camera. Seven days old females were weighed to the nearest μg. The number of eclosed, the number of stuck or dead pupae was quantified in each vial 15 days after cross initiation.

#### Visualization of acidic regions of the *Drosophila* gut

At least ten 10-day-old female flies were place on fly food supplemented with 2% bromophenol blue for 24 hours at 25°C. After dissection, intestines were immediately imaged to avoid diffusion of dye using a SMZ800N stereo microscope and a Digital Sight 1000 microscope camera.

#### Lysotracker staining

Ten female flies aged 3-10 days old were nutrient-depleted by placement in vials containing a cellulose plug infused with 3 mLs of PBS for 48 hours. At this point, intestines were removed and kept in ice cold PBS pending staining. Intestines were incubated for 5 minutes with 300 μl of a solution containing the Lysotracker probe (1:1000 dilution, Invitrogen) and DAPI (1 μg/ml) followed by three sets of 5 minutes washes in PBS. The samples were fixed in 4% formaldehyde for 20 minutes at room temperature and then washed three times. The guts were mounted in Vectashield mounting medium and analyzed using a confocal microscope, as described above.

#### Single cell RNA-seq

At least 200 midguts from each genotype were dissected under a dissecting microscope and placed in a dissection plate containing ice cold 1X PBS within a 2-hour time period. After cutting intestines into small pieces, they were centrifuged at 5,000 rpm for 5 minutes at 4 °C. Intestines were then placed in a 1.5 mL microcentrifuge tube solution containing 1X PBS/0.4 %BSA and elastase (1mg/mL, Sigma-Aldrich) for 30 minutes at 37°C with gentle rocking. The digestion reaction was stopped by placing the cells in 1X PBS/0.4 %BSA. The cell suspension was filtered through 70 μm and then 30 μm cell strainers and centrifuged at 5,000 rpm for 5 minutes at 4 °C. After discarding the supernatant, the pellet was suspended in 400 μL of cold 1X PBS with 1% BSA. Cells were assessed for viability and quantity using ViaStain AOPI Staining Solution (Revvity, CS2-0106) with the Cellometer K2 Fluorescent Cell Counter (Nexcelom Bioscience). Brightfield imaging and fluorescence measurements were used to assess cell viability.

Droplet-based microfluidic Chromium X (10X Genomics, Inc.) was used for generating single-cell Gel Beads-in-emulsions (GEMs) as per the Chromium Next GEM Single-Cell 5’ v2 - CG000331 Rev F (10X Genomics demonstrated protocol). A maximum of 10,000 intestine cells per sample were loaded into each channel. The Chromium Next GEM Single-Cell 5’ v2 - CG000331 Rev F 10X genomics workflow was used for all downstream steps to produce sequencing-ready libraries. The resulting libraries were normalized and pooled for sequencing on a NovaSeq system (Illumina).

Sequence demultiplexing, barcode processing, alignment, and gene counting was performed using 10X Genomics software CellRanger 7.1.0. Cellbender v. 0.2.0 was used for cell (https://github.com/broadinstitute/CellBender) identification and ambient RNA background correction.

### Quantification and statistical analysis

qRT-PCR and bulk RNA-seq were performed on biological triplicates. For qRT-PCR, statistical tests performed are indicated in the figure legends. For RNA-seq, DEseq was used to calculate fold change and statistical significance accounting for multiple comparisons. Thresholds of 2-fold change and adjusted p value of 0.05 or less were used to identify significant differences in expression. A Panther overrepresentation test was used to identify biological processes that were differentially represented between the two conditions ^53^. Numbers of prospero, tachykinin and labial positive cells were counted manually. For lysotracker quantification, Image J (FIJI) was used to quantify total fluorescence. This was normalized by dividing by the total area included in the measurement.

Background fluorescence was subtracted from this measurement. Gut acidity was quantified using Image J, as previously described ^79^. Data were graphed and appropriate statistical analyses were applied using Graphpad prism, version 11.

### Data availability statement

The bulk RNA-seq data has been deposited in the Geo Repository with accession number GSE330182. The single cell RNA-seq data has been deposited in the Geo Repository with accession number GSE325455.

---

**Fig S1:**
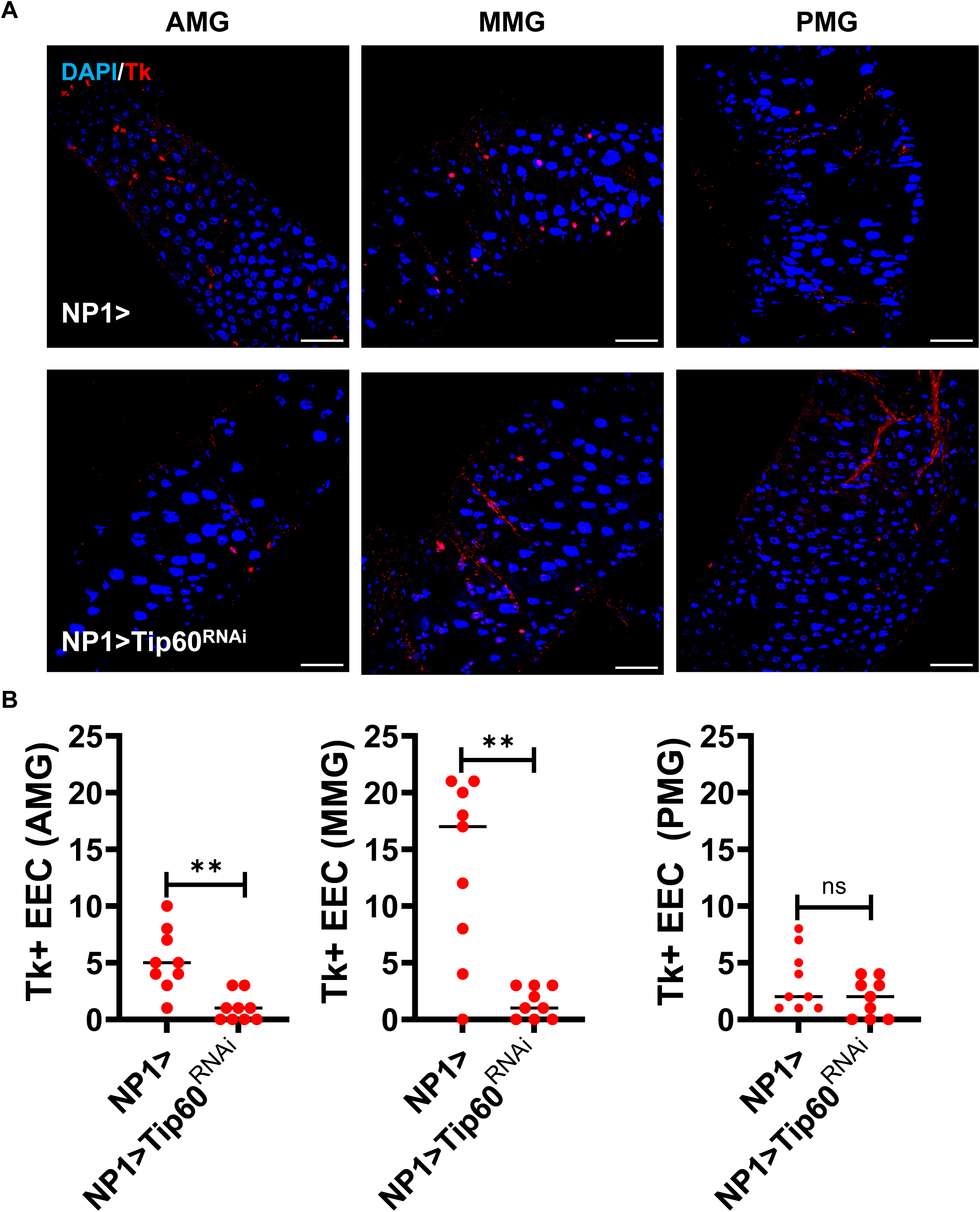
Fewer Tk+ enteroendocrine cells are observed in the AMG and MMG when Tip60^RNAi^ is expressed in enterocytes, related to Fig 1. (A) Tk immunofluorescence and (B) quantification of Tk+ enteroendocrine cells (EEC) in the AMG, MMG, and PMG. The mean of 9 intestines is shown. A Welch’s t test was used to assess significance. Measure bar 50 µM. ** p<0.01, ns not significant.

**Fig S2:**
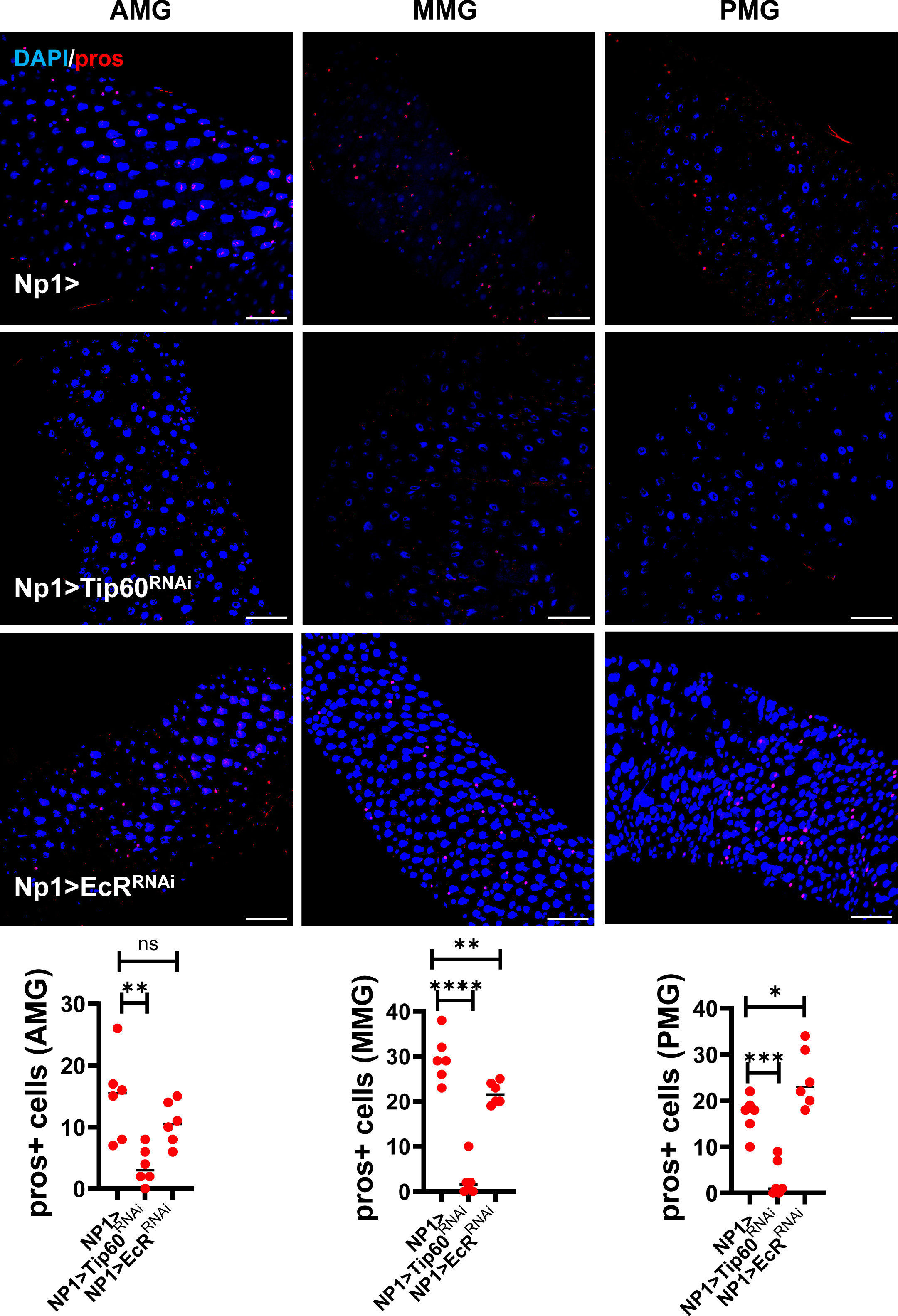
NP1>EcR^RNAi^ intestines have EEC numbers similar to those of NP1> flies, related to Fig 1. (A) pros immunofluorescence and (B) quantification of pros+ cells in the AMG, MMG, and PMG. The AMG data is also shown in Fig 1J. The mean of 6 intestines is shown. A one-way ordinary ANOVA with Dunnett’s multiple comparisons test was used to assess significance. Measure bar 50 µM. **** p<0.0001, *** p<0.001, ** p<0.01, * P<0.05, ns not significant.

**Fig S3:**
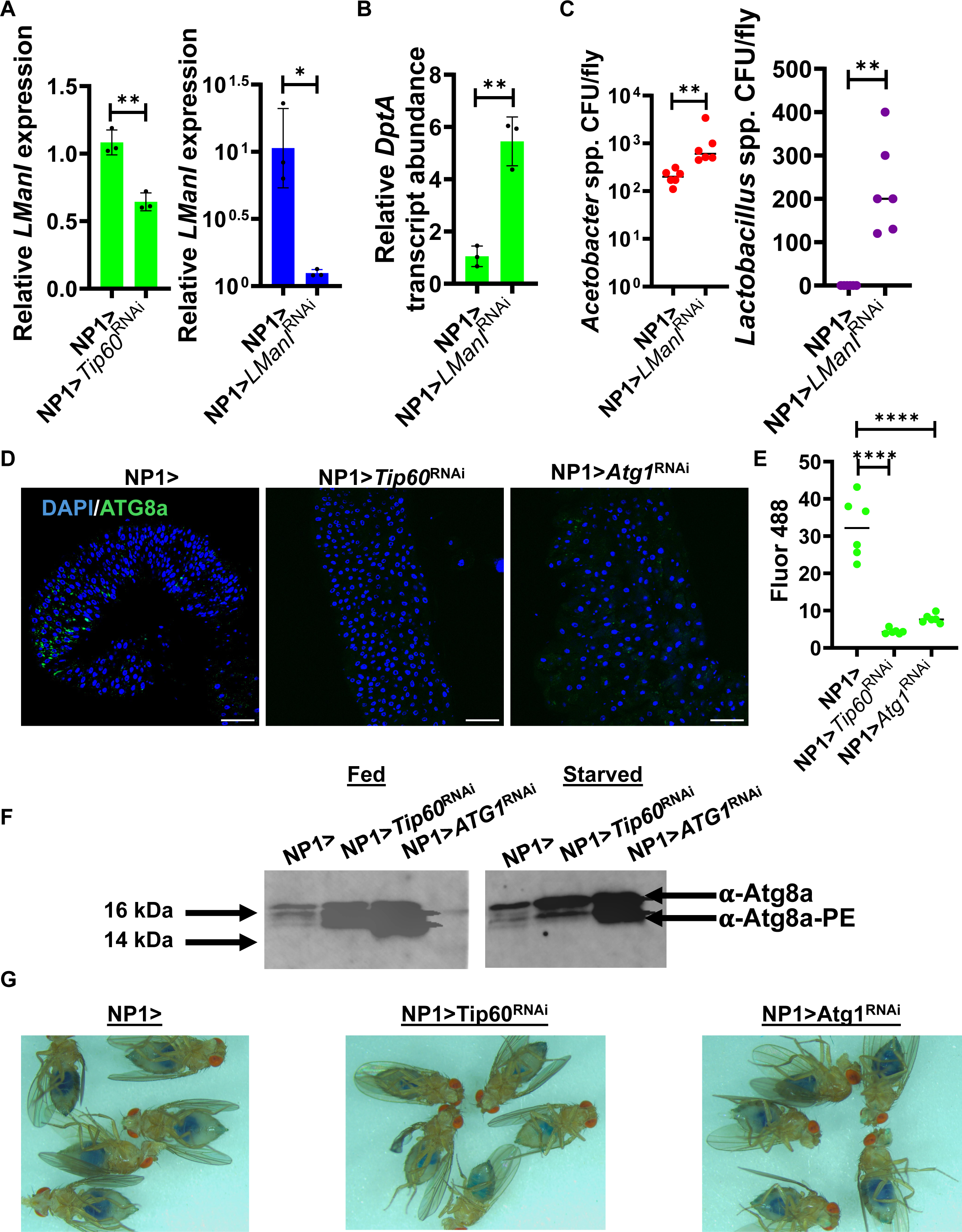
Evidence that Tip60RNAi in enterocytes blocks autophagy, related to Fig 3. (A and B) qRT-PCR of the indicated genes and fly genotypes. The mean of biological triplicates is shown. Error bars represent the standard deviation. An unpaired t test was used to determine significance of differences in *Lmanl* and *DptA* transcript abundance, while a Welch’s t test was used to determine differences in *Lmanl* transcript abundance. (C) Quantification of the burden of commensal bacteria in flies of the indicated genotypes. The mean of 6 flies is shown. A lognormal t test was used to determine significance of the *Acetobacter* measurement, while a Welch’s t test was used to determine the significance of the *Lactobacillus* measurement. (D) Micrographs and (E) quantification of ATG8a immunofluorescence in the intestines of starved flies of the indicated genotypes. The mean of 6 measurements is shown. A one-way ordinary ANOVA was used to determine significance. (F) Western blot analysis of unlipidated and lipidated Atg8a (Atg8a-PE) in whole fed or starved flies of the indicated genotypes. (G) SMURF assay of multiple flies of the indicated genotypes. **** p<0.0001, ** p<0.01, * p<0.05.

**Fig S4:**
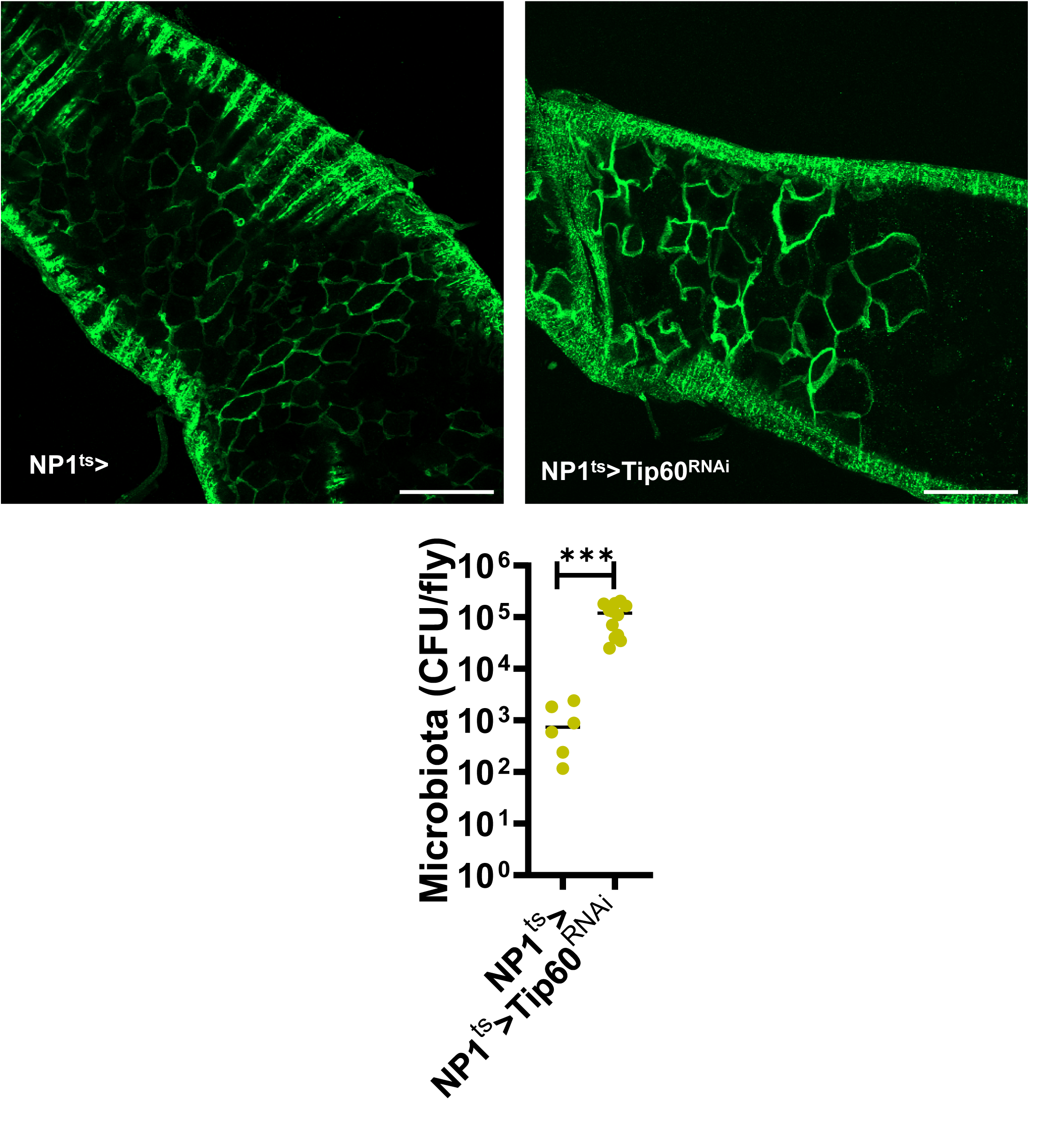
Microbial burden is elevated despite intact septate junctions, related to Fig 3. (A) Dlg immunofluorescent micrographs of the AMG of the indicated fly lines. A z projection is shown. Measure bar 50 µM. (B) Microbial burden of the same fly lines. *** p<0.001.

**Figure S5:**
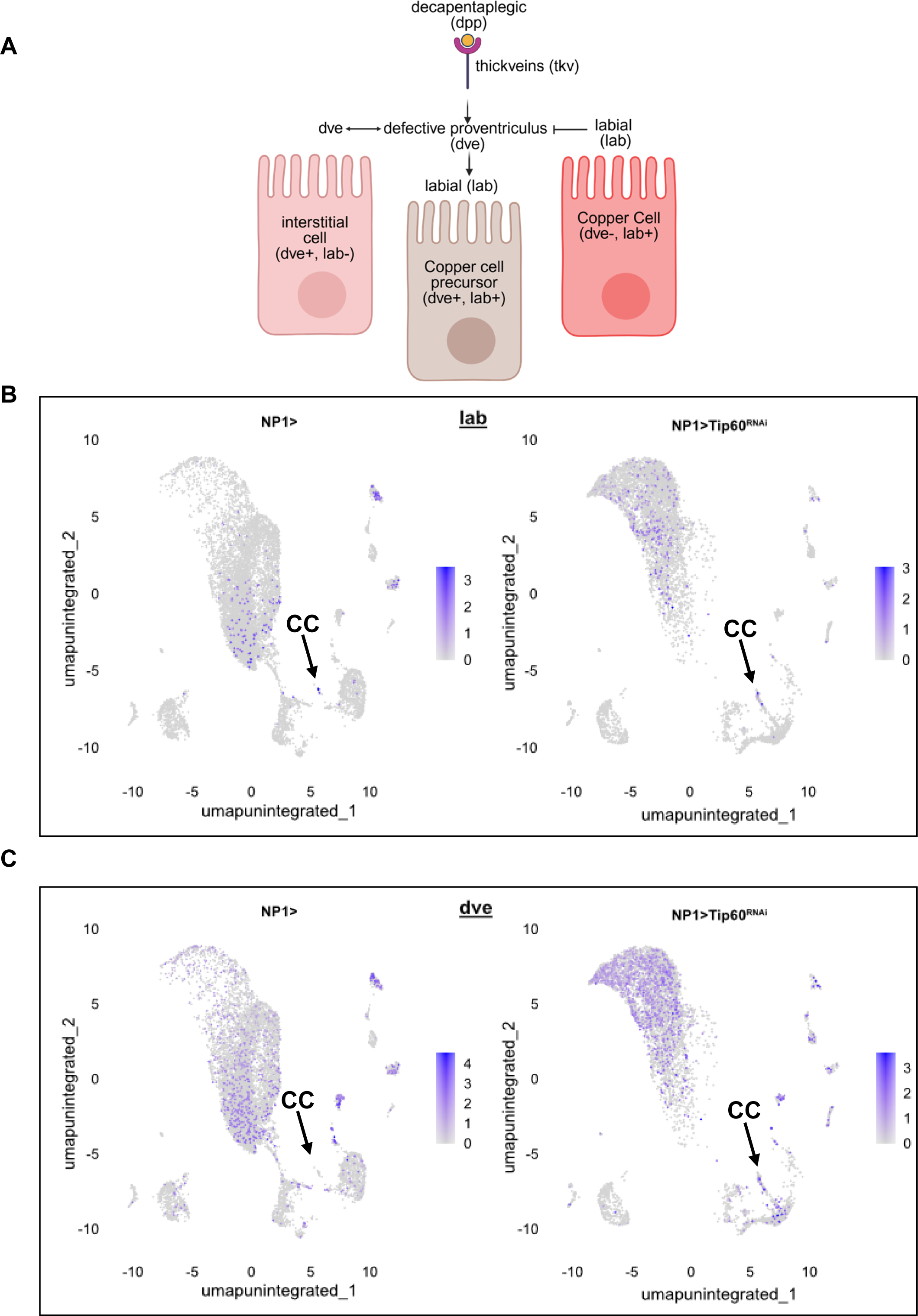
Evidence that the copper cells cluster of NP1>Tip60^RNAi^ flies is principally comprised of interstitial cells, related to Figure 7. (A) Schematic of the developmental program of the copper cell region of the MMG. Created in BioRender. Watnick, P. (2026) https://BioRender.com/d7w9auw. Umaps showing single cell expression of (B) lab and (D) dve in our data set. The copper cell region (CC) is indicated by an arrow.

**Figure S6.**
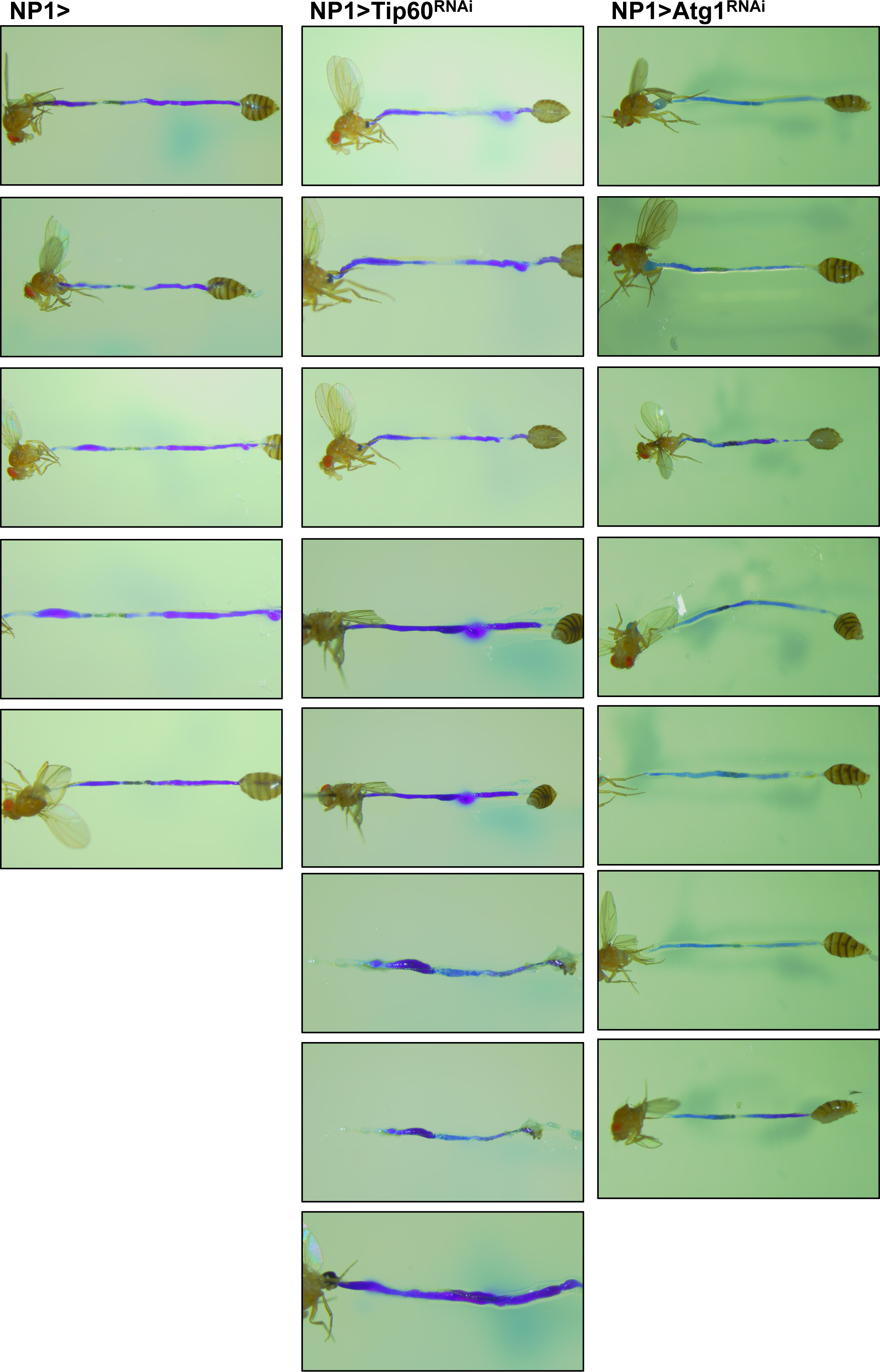
Bromophenol blue staining demonstrates decreased acid production in the intestines of NP1>Tip60^RNAi^ and NP1>Atg1^RNAi^ flies, related to Fig 7. Images of additional intestines of the indicated fly lines fed 0.1% bromophenol blue for 24 hours.

**Table S1:** Genes with increased transcript abundance in developing enterocytes that were shown to be highly expressed on in R3 in previous studies.

| Gene | Function | Log2 FC | Cell type (Cluster) |
| --- | --- | --- | --- |
| CG43774 | Unk | 6.23 | DE(6) |
| ArgK | Arginine kinase | 6.12 | DE(6) |
| CG5399 | Unk | 5.50 | DE(6), EB(1), EB(3), EB(4), EB(12), AMG(9) |
| pcl | Polycomb-like, midgut development | 4.95 | DE(6), ISC(0), EB(1), EB(4) |
| Tsp42Ec | Tetraspanin | 4.95 | DE(6), EB(4), LFC(10) |
| CG13323 | Unk | 4.90 | DE(6), EB(12), AMG(9) |
| CG34288 | Unk | 4.67 | DE(6) |
| CG43773 | Unk | 3.50 | DE(6) |
| CG1246 | Unk | 3.40 | DE(6) |
| CG15422 | Unk | 2.47 | DE(6), CC(8) |
| CG8051 | Predicted to be involved in monocarboxylic acid transport. | 2.45 | DE(6), EB (1) |
| CG15127 | Unk | 2.03 | DE(6) |
| CG15423 | Unk | 1.97 | DE(6), EB(3), CC(8) |
| Syx4 | superfamily of SNARE genes, which mediate membrane fusion. | 1.62 | DE(6) |
| hebe | determination of adult lifespan | 1.54 | DE(6), ISC(0), ISC(8), EB(1), EB(4) |

